# Single pulse electrical stimulation in white matter modulates iEEG visual responses in human early visual cortex

**DOI:** 10.1101/2025.05.05.652264

**Authors:** Harvey Huang, Kendrick N. Kay, Nicholas M. Gregg, Gabriela Ojeda Valencia, Myung-Ho In, Christoph Kapeller, Yunhong Shu, Gregory A. Worrell, Kai J. Miller, Dora Hermes

## Abstract

Electrical stimulation is increasingly used to modulate brain networks for clinical purposes. The basic unit of neurostimulation, a single electrical pulse, can travel through white matter to influence connected neuronal populations. However, the mechanisms by which it influences connected populations is not well understood: stimulation may excite, inhibit, or add noise to neuronal population activity. In this study, we investigated how single pulses modulate the neuronal processing of images in a well-controlled visual paradigm. In two human subjects implanted with iEEG electrodes for clinical purposes, single pulses were delivered to electrodes in white matter tracts connected to measurement electrodes in visual cortex. Images appeared on-screen at 0, 100, or 200 ms after each pulse. Using finite impulse response modeling, we decomposed the broadband and evoked potential responses into separate components induced by electrical stimulation and by visual processing. Single pulses induced transient broadband increases followed by suppression, but they did not modulate the visual broadband responses (i.e., stimulation response was additive to visual response). In contrast, single pulses elicited prominent brain stimulation evoked potentials *and* they modulated the visual evoked potentials. Specifically, visual evoked potentials were larger when stimulation occurred closer to visual onset. This indicates that a single electrical pulse can increase the strength or synchrony of visual inputs. Overall, these findings suggest that the effects of electrical stimulation in the visual system are two-fold: stimulation induces additive effects on broadband power, possibly by adding noise, and it interacts with synchronous visual inputs to amplify them.

## 1. Introduction

Intracranial neurostimulation has shown great promise as a treatment for a variety of neurological conditions. Current FDA-approved neurostimulation therapies reduce symptom burden for patients with movement disorders and medication-refractory epilepsy^1–3^. Emergent neurostimulation efforts also exist to restore vision^4–6^ and treat psychiatric disorders such as depression and obsessive-compulsive disorder^7–9^. Many such applications target well-connected brain nuclei (e.g., thalamus) or white matter, and are thought to function by affecting connected areas within brain networks^10^. Despite clinical success, the mechanisms by which neurostimulation modulates natural dynamics within brain networks remain the subject of ongoing investigation, with some theories suggesting that stimulation virtually lesions neuronal populations^10,11^, and others suggesting that stimulation injects neuronal noise^12,13^. Virtual lesions would be expected to suppress ongoing neuronal processing, whereas injected neuronal noise might result in added neuronal activity or nonlinear effects such as stochastic resonance^14,15^. These theories can be tested by carefully characterizing the interaction between electrical stimulation and neuronal activity.

A single electrical pulse is the basic unit of most neurostimulation therapies and is sufficient, in itself, to produce dynamic effects on local neuronal activity at connected measurement sites^16^. For example, single pulse electrical stimulation (SPES) through microelectrodes in macaque thalamus can elicit a transient increase followed by a sustained decrease in neuronal firing in connected visual cortex^17^. This resembles our previous population-level intracranial EEG (iEEG) findings, where single electrical pulses delivered to the hippocampus induced transient increases followed by sustained decreases in broadband power in the ventral temporal cortex^18^. These broadband power changes are thought to correlate with neuronal population firing rates^19–21^ and represent a stimulation effect on population activity.

Whether SPES modulates neuronal activity not just at rest but also during active engagement remains a gap in understanding. We address this in the present study by combining iEEG SPES with a visual task in two human participants. We stimulated major white matter tracts connected to early visual areas (V1-V3) while presenting a visual stimulus at three different intervals after electrical stimulation. We separated the iEEG responses into stimulation-driven and visual-driven components and tested whether the visual components differed by stimulation condition. If so, this was taken as evidence that SPES modulates visual processing. We quantified stimulation-visual interactions for both broadband power changes and evoked potentials, which are two independent iEEG signal features that reflect complementary neurophysiological events^22,23^. Broadband power is related to asynchronous population firing^19– 21^, while evoked potentials quantify synchronous synaptic inputs^24,25^. Interpreting stimulation effects on broadband power and evoked potentials together helps us advance toward a comprehensive picture of how electrical pulses interact with dynamic neuronal activity.

## 2. Methods

### 2.1. Subjects and iEEG recording

iEEG data were recorded in 2 human subjects (subject 1: 21M, subject 2: 18M) who had been implanted with stereo-EEG electrodes for epilepsy seizure localization. Subjects provided informed consent and the study was conducted according to the guidelines of the Declaration of Helsinki and approved by the Institutional Review Board of the Mayo Clinic (IRB #15-006530), which also authorizes sharing of deidentified data. In each subject, iEEG electrodes were placed in the visual cortex and in other brain areas (subject 1: 192 electrodes, subject 2: 238 electrodes), as dictated by the clinical team. Recorded data were digitized at 4800 Hz on a g.Hiamp biosignal amplifier (g.tec medical engineering GmbH., Austria), and originally referenced to an electrode in the white matter.

### 2.2. Task and stimuli

The main experiment combined SPES with a visual perceptual decision-making task (referred to as SPES+Visual). During the task, subjects fixated at the center of the screen, spanning ∼28 degrees of visual angle at ∼60 cm distance. Grayscale images constructed from two natural scenes were presented for 1 s each with 1-1.4 s of rest (uniform gray screen) in between (Figure 1A), via custom g.HIsys Simulink software (g.tec). Participants were instructed to press a button to indicate which of two natural scenes was reflected in each image presented. Subject 1 also received short auditory feedback upon each button press to indicate response correctness (a “coin jingle” when correct, a buzz when incorrect).

**Figure 1.**
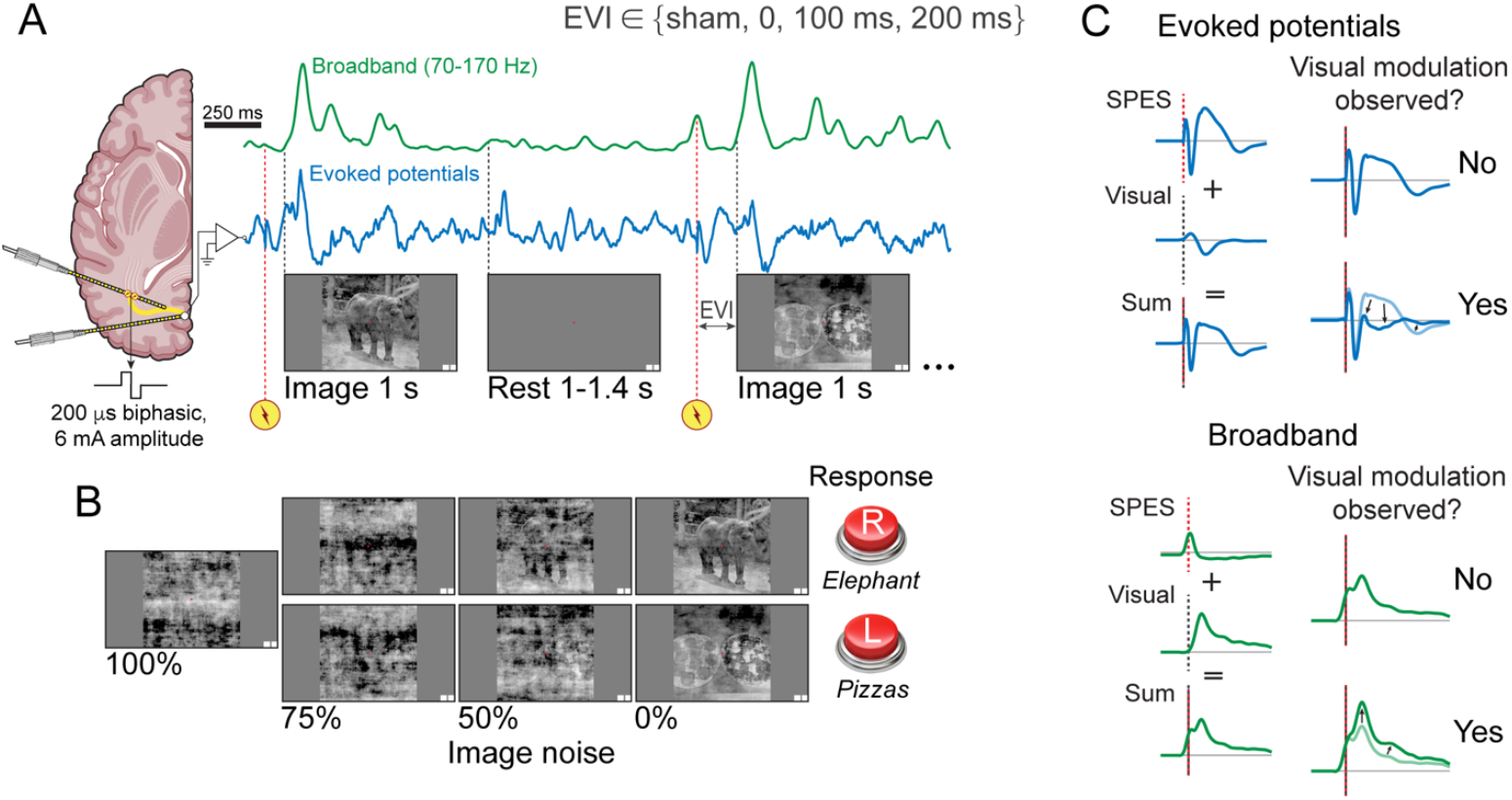
SPES+Visual experimental task and objectives. **A**, Subjects fixated on images of 1 s duration with 1-1.4 s rest, and SPES occurred in non-sham trials at 0, 100, or 200 ms before visual onset (EVI). Evoked potentials were recorded and transformed to time-varying broadband estimates. Brain graphic created in part with BioRender.com. **B**, Seven image conditions corresponded to two natural scenes mixed with four levels of noise. Subjects were instructed to identify the natural scene by button press. **C**, The primary objective was to determine whether visual evoked potentials and visual broadband changes were modulated by SPES.

The images were produced by modifying the phase spectra of two natural scenes, *Elephant* and *Pizzas*, according to the procedure in Heekeren et al. (Figure 1B).^26^ This procedure generated seven unique image conditions: *Elephant* or *Pizzas* mixed with 0%, 50%, or 75% noise, and a pooled 100% noise condition. First, each natural scene was transformed into the frequency domain using a 2D fast Fourier transform (FFT), producing a magnitude matrix and a phase matrix. The magnitude matrices from the two natural scenes were averaged to create a common magnitude matrix, ***M***. Then to create images with different levels of noise, the phase matrix from either *Elephant* or *Pizzas* was mixed with a random noise matrix, whose values were drawn from a uniform distribution between 0 and 2*pi, at mixing ratios of 100:0 (original phase), 50:50, 25:75, or 0:100 (entirely noise). Each mixed phase matrix was combined with ***M***, and the final image was reconstructed using an inverse 2D FFT. Correctness in the 100% noise condition was still coded by the original natural scene used to create each image.

SPES (single biphasic pulses, 100 μs per phase, 6 mA) was delivered to target stimulation sites before the onset of each image (Figures 2, S1). The electrical-visual time interval (EVI) between SPES and visual image onset was 200 ms, 100 ms, or 0 ms. Some trials included no stimulation (“sham” condition). The experiment was organized into 7-minute runs. Subject 1 performed 2 runs in one session and 2 additional runs 4 days later. Subject 2 performed 3 total runs in a single session.

**Figure 2.**
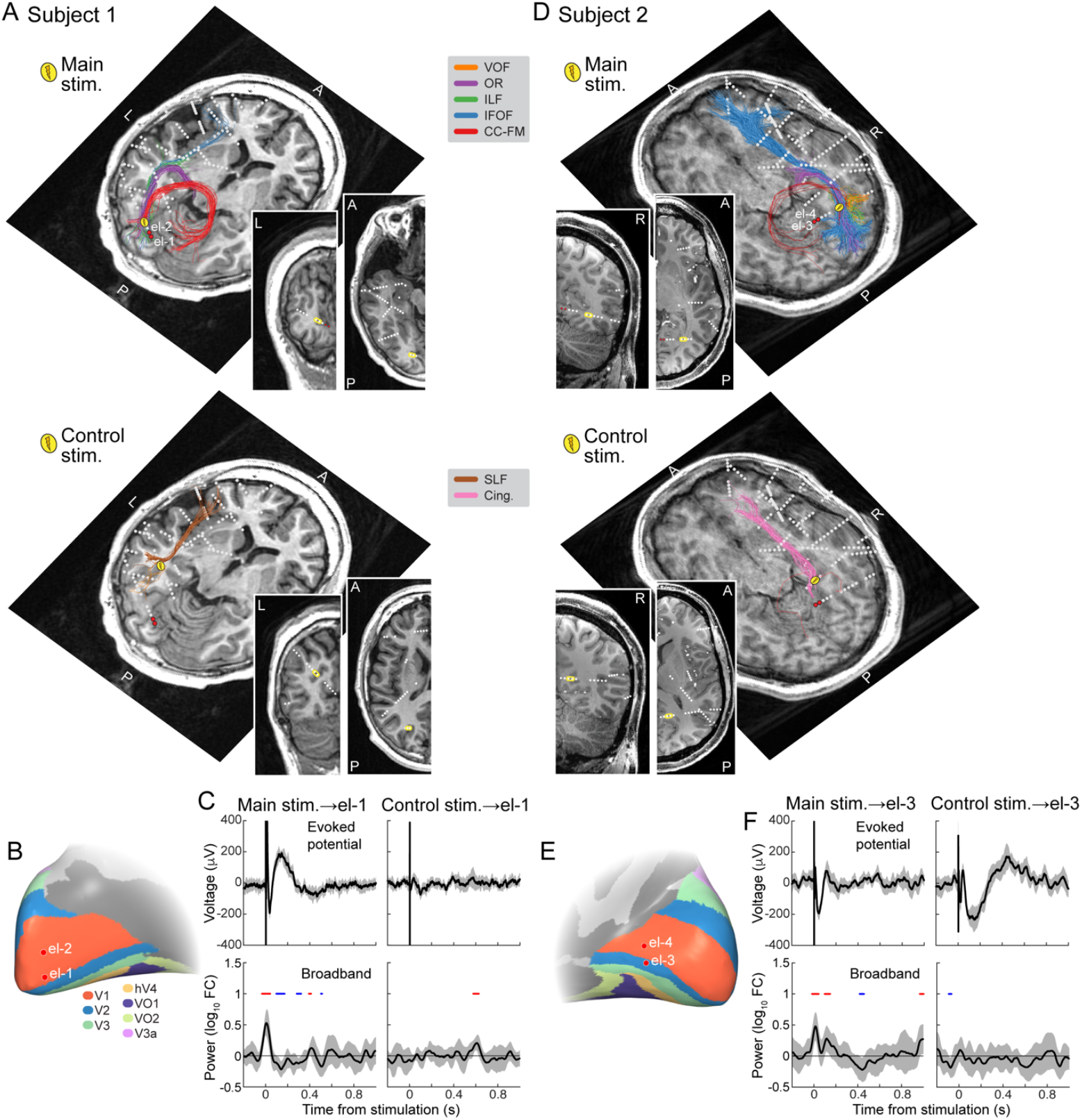
Stimulation sites and EVC measurement electrodes. **A**, Main and control stimulation sites, and early visual measurement electrodes (el-1 and el-2) in subject 1, plotted alongside white matter tracts within 4 mm of each stimulation site. VOF = vertical occipital fasciculus, OR = optic radiation, ILF = inferior longitudinal fasciculus, IFOF = inferior fronto-occipital fasciculus, CC-FM = corpus callosum forceps major, SLF = superior longitudinal fasciculus, Cing. = cingulum. Insets show coronal and axial T1-weighted MRI slices centered on each stimulation site. **B**, Early visual measurement electrodes on the subject’s inflated occipital pial surface. **C**, BSEPs (top) and stimulation-induced broadband changes (bottom) measured at el-1 from each stimulation site. Shaded intervals depict 95% confidence interval of the mean. Time points with mean broadband significantly greater than or less than 0 are highlighted in red and blue, respectively (one-sample *t*-test, P < 0.05). **D-F**, Stimulation sites and early visual measurement electrodes (el-3 and el-4) in subject 2, as in A-C. BSEPs and stimulation-induced broadband changes measured at el-2 and el-4 are shown in Figure S1.

The 7 image conditions and 4 stimulation conditions (3 EVIs + sham) yielded 28 unique experimental conditions per stimulation site. There were 6-24 trials per experimental condition, yielding 216-384 total trials for each stimulation site.

### 2.3. Independent SPES data

To understand the brain’s responses to electrical stimulation alone, we first delivered SPES separately without visual stimuli, while each subject lay awake at rest (Figure 2C, F). This occurred immediately after the first SPES+Visual session for subject 1 and immediately before the session for subject 2. At each stimulation site, SPES was delivered 12 times (trials) with 2-5 second intervals between pulses, using the same stimulation parameters as in the SPES+Visual experiment.

### 2.4. Selection of measurement and stimulation electrodes

Electrode locations were identified from the postoperative CT scan and coregistered to the preoperative T1-weighted MRI^27^, which was transformed into AC-PC space by affine transformations and trilinear voxel interpolation^28^. Measurement electrodes in early visual cortex (EVC: V1, V2, V3) were anatomically identified (numbered 1-4, Figure 2) based on automated segmentation of subject T1-weighted MRIs using Freesurfer 7^29^, the Benson probabilistic visual atlas, and the Wang atlas^30,31^. Our primary analysis focused on these EVC electrodes.

Additional findings from visually responsive electrodes outside of the EVC (identified functionally, see supplemental methods) are presented separately (in section 3.4 and Figures S4, S5).

We stimulated electrode pairs located near major white matter tracts. The proximity of all electrodes to white matter tracts was estimated based on diffusion MRI (see supplemental methods). Whole brain tractography was performed using probabilistic tracking with constrained spherical deconvolution. From the whole brain connectome, fiber bundles were recognized using the pyAFQ^32^ package with RecoBundles^33^ and the 80 bundle HCP atlas^34^. We estimated how many fiber streamlines passed within 4 mm of each stimulation electrode and how many endpoints landed within 6 mm of each EVC electrode, for the following white matter tracts of interest: VOF = vertical occipital fasciculus, OR = optic radiation, ILF = inferior longitudinal fasciculus, IFOF = inferior fronto-occipital fasciculus, CC-FM = corpus callosum forceps major, SLF = superior longitudinal fasciculus, Cing. = cingulum. In each subject, the “main” stimulation site was located near a dense cluster of streamlines in the IFOF and ILF, which had endpoints in EVC. Stimulation sites located farther from the IFOF and ILF were selected as “control”.

### 2.5. iEEG data preprocessing

First, after visual inspection of all data, electrodes were excluded if they contained electrophysiological artifacts, were located outside of brain tissue, or were part of seizure onset zones per physician records. Individual trials containing epileptiform activity were also removed. Second, data were high-pass filtered (2^nd^ order Butterworth with cutoff frequency = 0.3 Hz, applied in a forward-reverse manner) to remove low-frequency drift. Third, data were re-referenced to an adjusted common average reference, calculated using the least responsive 25% of electrodes (based on activity 10-300 ms after visual onset) to minimize the introduction of large responses during re-referencing^35^. Finally, line noise at 60 Hz and its first two harmonics (120 and 180 Hz) were attenuated using a spectrum interpolation method^36^, resulting in clean evoked potential data.

Time-varying broadband responses (Figure 1A, green) were calculated from the clean evoked potential data as follows. Transient stimulation artifacts from SPES were removed by linear interpolation between 0 and 4 ms after stimulation. For each experimental condition (image × EVI), the mean evoked potential across trials was subtracted to minimize filtering artifacts due to sharp evoked potential peaks. Each trial was forward-reverse filtered with third order Butterworth bandpass filters, divided into 20 Hz-wide bands between 70 and 170 Hz, excluding 110-130 Hz to avoid contamination by residual line noise harmonics. Power within each band was quantified as the squared envelope of the analytic signal (absolute value of the Hilbert transform); and the overall broadband power was the geometric mean across all bands. The broadband power was normalized by baseline, which was defined as the geometric mean broadband power across all sham stimulation trials 250-50 ms before visual onset, calculated separately for each run. Final normalized units for broadband power were expressed as “fold increase” from baseline after subtracting 1. For example, this results in a 0.5-fold increase for a time point with broadband power = 9 μV^2^ and baseline = 6 μV^2^ (9 ÷ 6 ™ 1 = 0.5). As the choice of iEEG reference may impact the estimation of broadband, we repeated broadband calculations and its associated analyses on bipolar re-referenced data, where the adjusted common average referencing step was replaced by the difference in evoked potentials between adjacent measurement electrodes (see supplemental results).

### 2.6. Finite impulse response analysis of stimulation and visual responses

Since stimulation occurs shortly before visual onset in each trial, the stimulation and visual responses at least partially overlap in time. To separate the stimulation and visual component responses, we used finite impulse response (FIR) modeling, first for evoked potentials and then for broadband responses. These models test whether the observed responses are better explained by visual component responses that remain unchanged with stimulation (reflecting a pure additive relationship between stimulation and visual responses) or by visual component responses that change with varying EVI (reflecting an interaction between stimulation and visual responses).

Here, we describe the FIR analysis for evoked potentials, which includes FIR modeling, model comparison, and final model selection steps. The FIR analysis proceeded similarly for broadband responses (see supplemental methods for details). First, evoked potentials were downsampled to 600 Hz. We considered four FIR models of the linear form y = Xβ, where y represents the vector of observed evoked potentials concatenated across all trials, X is the design matrix with finite impulse response predictors for hypothesized component evoked potentials across time points, and β consists of the voltage coefficients for each component across time points, estimated via least squares regression (Figure 3). The first and second models assume that all observed evoked potentials are the sum of a brain stimulation evoked potential (BSEP) and visual evoked potentials (VEPs) that do not change with stimulation:

**Figure 3.**
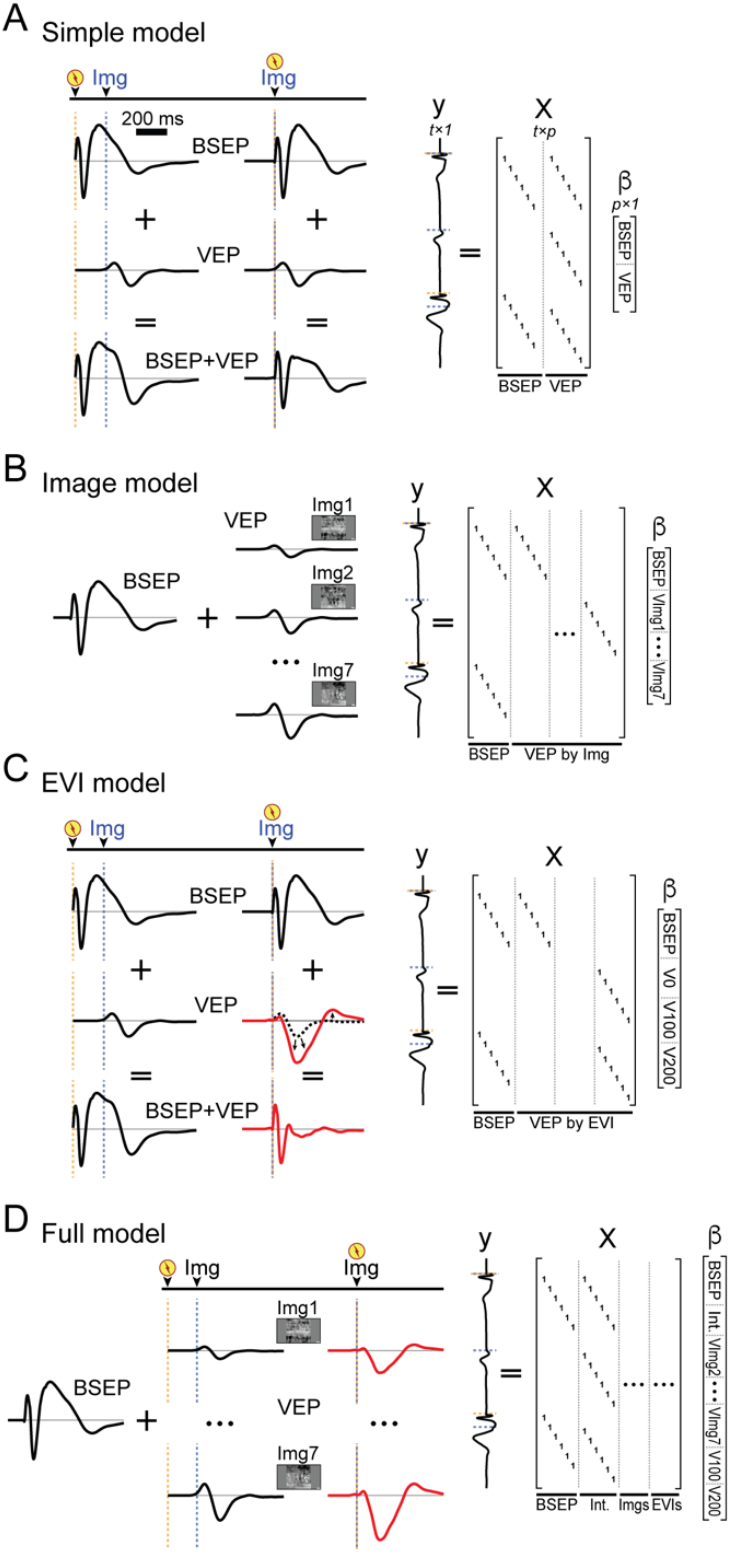
FIR models fit on SPES+Visual evoked potentials. **A**, The simple model fits a single set of BSEP predictors and a single set of VEP predictors, regardless of image condition or EVI. **B**, The image model fits independent VEP predictors for each of the seven image conditions. **C**, The EVI model fits independent VEP predictors for each of the three EVI conditions (sham is merged with 200 ms). **D**, The full model fits nine independent sets of VEP predictors, corresponding to all image and EVI conditions.

1. <u>The simple model</u> contains two sets of predictors: one for the BSEP 0-1 s post-stimulation, and one for the VEP 0-1 s after visual onset, regardless of EVI or image identity (Figure 3A).
2. <u>The image model</u> contains eight sets of predictors: one for the BSEP, and a set of VEP predictors for each image condition, which allow VEPs to vary by image (Figure 3B). The third and fourth models allow visual responses to vary by stimulation condition:
3. <u>The EVI model</u> contains four sets of predictors: one for the BSEP, and a set of VEP predictors for each EVI (0 ms, 100 ms, and 200 ms) (Figure 3C). Sham stimulation trials were assigned the same VEP predictors as the 200 ms EVI trials, by model necessity.
4. <u>The full model</u> contains ten sets of predictors: one for the BSEP, and nine sets of VEP predictors that vary by both image condition and EVI in a two-way design. The first set of VEP predictors can be thought of as an “intercept” condition for all trials with 100% image noise and 0 ms EVI; the next six capture the difference from all other image conditions to the intercept; and the last two capture the difference from all other EVIs to the intercept.

The four FIR models were compared, separately for each stimulation–measurement electrode pair. To control for the different numbers of free parameters in the models, we used split-half validation to quantify model accuracy. Odd-numbered trials within each experimental condition were pooled as training data for each FIR model, and the even-numbered trials were held out for testing. Model performance was assessed using the coefficients of determination (COD):

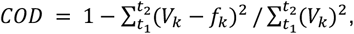

where *V*_*k*_ is the observed voltage at time point *k, f*_*k*_ is the predicted voltage at time point *k, t*_*1*_ is the earliest predictor time point (different for each trial), and *t*_*2*_ is fixed at 0.5 s after visual onset, which approximated the end of most meaningful evoked potentials.

The models we designed are naturally ordered in terms of complexity, and we used split-half validated COD to determine whether any of the more complex models are necessary to explain the data. Whenever the simple model had the highest mean COD or when no other model was significantly better, we concluded that the simple model fit best. Whenever either the image or the EVI model significantly outperformed the simple model (by paired *t*-test on COD, right-tailed P < 0.05), we concluded this to be the best model. Finally, whenever the full model significantly outperformed the next best model, we concluded that the full model fit best. As a baseline comparison, COD was also computed directly from the data itself, where the prediction for each testing trial was simply the mean across training trials of the same experimental condition.

We checked whether button presses influenced evoked potentials using additional FIR predictors, but did not find that such a model explained additional variance. We calculated bootstrapped confidence intervals for all FIR voltage coefficients (β). These steps are described in the supplemental methods.

### 2.7. Psychometric analysis

We recorded reaction times and accuracies for each subject. Trials without responses were omitted. To examine whether reaction times differed across the seven image conditions, we performed a one-way ANOVA. Additionally, a multivariate fixed effects linear regression model was created to evaluate whether reaction time varied with run number, trial onset time, image condition (categorical), and stimulation condition (categorical). Stimulation condition had seven levels: sham, and stimulation at either main or control stimulation site at each EVI (0, 100, 200 ms).

For each image condition, a right-tailed binomial test determined whether response accuracy was significantly higher than expected by chance (50%). For each stimulation condition, we modeled response accuracy as a function of image noise using the best-fit Weibull function:

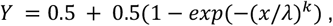

where *Y* is the response accuracy between 0 and 1, *x* is the image coherence (the complement of image noise, 100% − image noise), and *λ* and *k* are parameters. For this analysis, trials were pooled across *Elephant* and *Pizzas* images of the same image noise level, resulting in four distinct levels in the independent variable (100%, 75%, 50%, or 0% image noise). For levels of 75%, 50%, and 0% image noise, we explicitly tested whether response accuracy depended on stimulation condition using Chi-Square tests of independence. We also fit two multivariate logistic regression models to test whether response accuracy varied first with image coherence, original natural scenes identity, and their interaction; and second with image coherence, run number, trial onset time, and stimulation condition.

## 3. Results

To understand how single pulse electrical stimulation (SPES) affects ongoing neural processing, we stimulated electrodes in the white matter projecting to early visual cortex in two human subjects. The degree of interaction between stimulation and visually driven iEEG responses in the early visual cortex was assessed in order to test whether stimulation modulates image processing. We assessed two iEEG signal features that provide complementary information about neuronal activity: evoked potentials and broadband power changes.

### 3.1. Stimulation of visual white matter pathways produces evoked potentials and broadband power changes at early visual electrodes

We first stimulated visual and control pathways in the absence of any task to examine the neurophysiologic effects produced by stimulation alone. SPES was delivered at two stimulation sites in each subject: a main stimulation site near major visual pathways (<4 mm from the OR, VOF, ILF, IFOF, and CC-FM), and a control stimulation site located near the SLF and cingulum bundles. Measurement electrodes were located in V1-V2 (electrodes 1-2 in subject 1, Figure 2A-B, electrodes 3-4 in subject 2, Figure 2D-E). Diffusion MRI confirmed that streamlines connect measurement electrodes to the main stimulation site, but not to the control stimulation site. Specifically, of all streamlines near the main stimulation sites, 23 (17%), 17 (12%), 8 (1.3%), and 11 (1.8%) passed within 6 mm of measurement electrodes 1-4, respectively.

Brain stimulation evoked potentials (BSEPs) did not correlate perfectly with anatomical connectivity, as stimulation of the main sites in both subjects as well as the control site in subject 2 produced prominent BSEPs in EVC (Figures 2C and F, S1, top). In contrast, stimulation-induced broadband changes did correlate with anatomical connectivity, as they were observed in EVC only when stimulating the main sites (Figures 2C and F, S1, bottom). These broadband changes were characterized by transient power increase within 50 ms of stimulation, followed by a return to baseline and then relative suppression lasting up to 400 ms in subject 1. This pattern of broadband response resembles what we previously observed with SPES^18^, and demonstrates that stimulation affects the local neuronal population activity selectively at connected sites.

### 3.2. SPES in visual pathways modulates visual evoked potentials

The effects of single electrical pulses on visual processing were assessed by the SPES+Visual task, in which the white matter pathways were stimulated either immediately before or concurrently as subjects saw grayscale images with varying levels of noise (Figure 1A, B). SPES was delivered at electric-visual time intervals (EVI) of 200 ms, 100 ms, and 0 ms, except in interleaved sham stimulation trials. We used FIR models to separate brain stimulation (BSEP) and visual (VEP) evoked potential components from the overall observed responses (Figures 4B-E). The BSEP components resembled the BSEPs recorded in the independent SPES task (Figure 2C), but with slightly greater amplitudes. Separately, visual images elicited robust VEPs in early visual cortex when no stimulation was delivered (“sham”, e.g., subject 1: Figure 4A). Then, to determine whether these VEPs were modulated by SPES, we examine whether the VEPs differed significantly by EVI.

**Figure 4.**
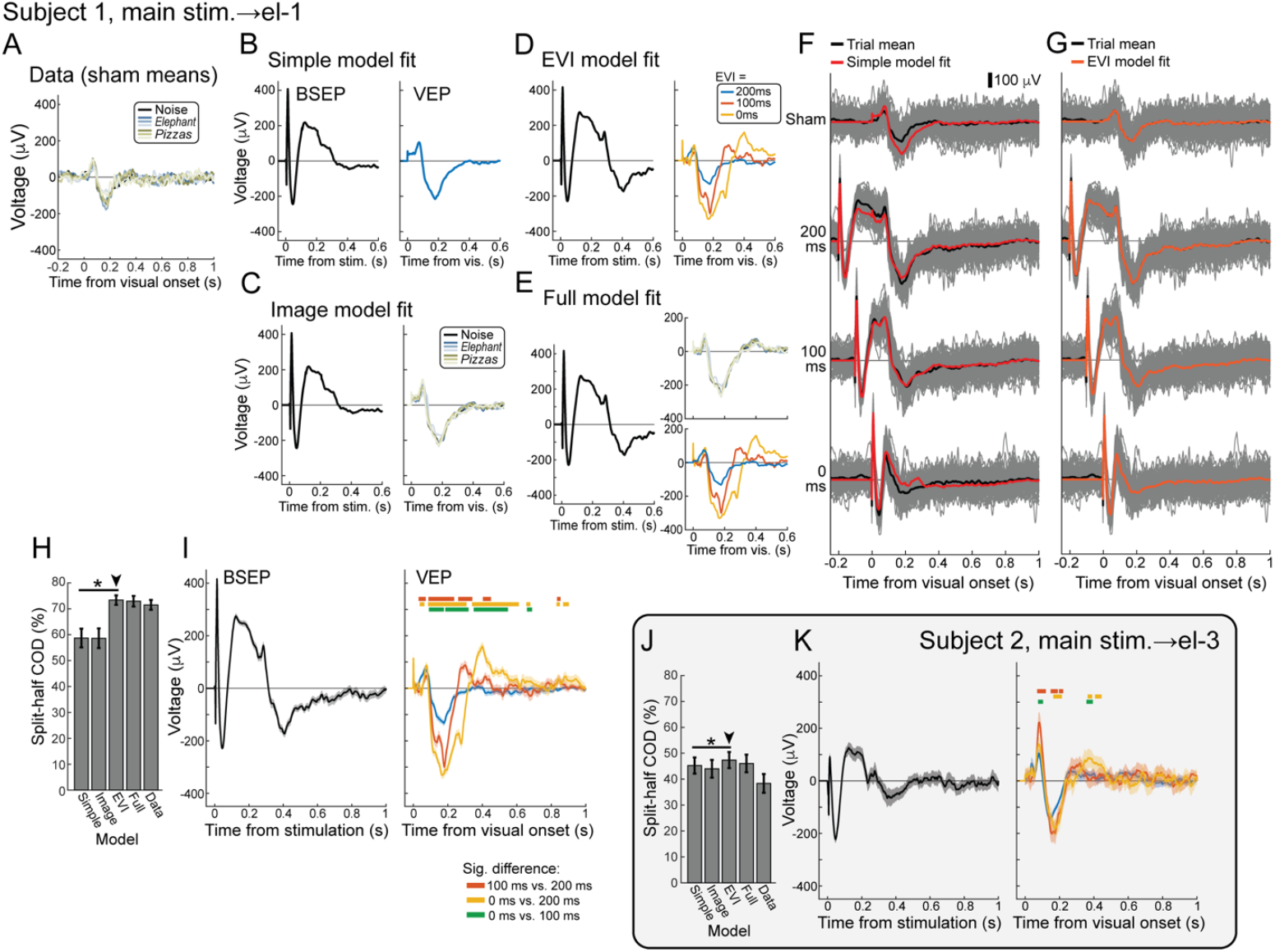
SPES in visual white matter tracts modulates VEPs at EVC electrodes. **A**, Mean of VEPs across sham stimulation trials at el-1, for each image condition. **B-E**, Component responses from simple, EVI, image, and full models fit on evoked potential data at el-1 from the main stimulation site in visual white matter. In the image model legend, lighter color denotes lower image noise (75%, 50%, 0%). **F**, Comparison between the data and the predictions from the simple model (B) at each EVI. All image conditions are pooled for each row. **G**, Comparison between the data and predictions from the EVI model (D) at each EVI. **H**, COD (mean and standard error) from testing trials for models in B-E. “Data” predicts test trials using the mean of condition-matched training trials. The EVI model yielded the highest COD, significantly greater than the simple model (*paired *t*-test, right-tailed P < 0.05) **I**, Bootstrapped mean and 95% confidence intervals for EVI model responses. Significantly different time intervals between pairs of VEPs fit for each EVI are labeled with colored bars (Bootstrapped Differences, P < 0.01). **J, K**, COD and EVI model responses (mean and 95% confidence interval) for evoked potential data at analogous main stimulation site to measurement electrode el-3 in subject 2. COD and EVI model responses for el-2 and el-4, as well as for the control stimulation sites, are presented in Figure S2.

The models show that VEPs differed significantly by EVI when visual white matter pathways (main stimulation sites) were stimulated (subject 1: Figure 4D, E, I, subject 2: 4K), but not when the control sites were stimulated (Figure S2B). When visual pathways were stimulated, the EVI model (which allows the visual response to vary with EVI) explained a significantly greater fraction of variance in the observed data (subject 1: COD = 73.4%, subject 2: COD = 47.4%, Figure 4H, J) than the simple model. The evoked potentials predicted by the EVI model (Figure 4G) indeed align better with observed data than those predicted by the simple model (Figure 4F). In contrast, VEPs did not vary by image noise level, as neither the image model nor the full model outperformed the EVI model. In all cases, the best-fit model also explained more variance in the data than unmodeled predictions calculated from the means of condition-matched training trials (“data”) (Figure 4H, J). In contrast, suboptimal models sometimes performed worse than data (e.g., simple and image models in 4H). In summary, the models demonstrate that a single electrical pulse delivered at or before visual image onset can rapidly modulate VEPs in the EVC.

How exactly were the VEPs modulated? In both subjects, the prominent negative peak at ∼180 ms after visual onset were larger when stimulation was delivered at or briefly before visual onset (EVI = 100 ms and EVI = 0 ms), compared to when stimulation was delivered 200 ms before visual onset or when there was no stimulation (Figure 4I, K). In subject 2, we note that stimulation at 100 ms before visual onset also significantly enhanced the amplitude of the initial positive VEP peak (at ∼100 ms after visual onset, Figure 4K).

### 3.3. SPES in visual pathways suppresses broadband power without modulation of visual responses

Whereas evoked potentials reflect synchronous synaptic inputs, induced broadband power changes capture asynchronous changes in local neuronal input firing rates^19–21^. We tested whether SPES also modulates visual broadband responses by comparing the same four FIR models applied to the observed broadband power changes.

SPES induced similar broadband responses during the visual task (captured by the FIR stimulation component) as SPES during rest. For both subjects, stimulation in visual pathways produced a transient increase in broadband power, followed by a decrease below baseline lasting ∼0.4 s in the EVC (Figure 5B, E).

**Figure 5.**
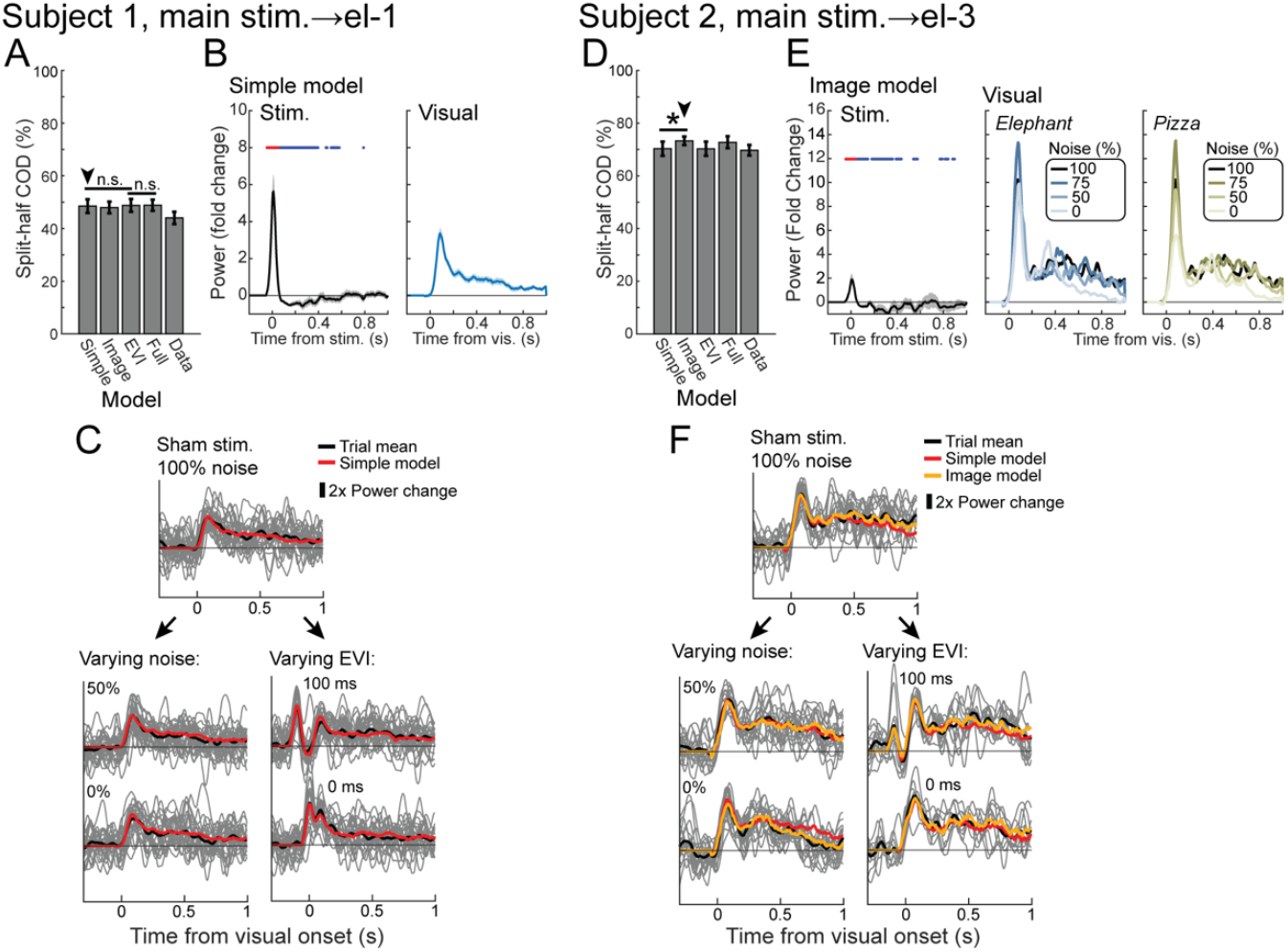
Independent stimulation-induced and visual induced broadband power changes at EVC electrodes. **A**, COD (mean and standard error) from testing trials for each model fit on broadband power at el-1 from main site stimulation. “Data” predicts test trials using the mean of condition-matched training trials. “n.s.” = not significant by paired *t-*test, right-tailed P > 0.05 (Full > EVI, EVI > Simple). **B**, Bootstrapped mean and 95% confidence interval for simple model responses. Time points significantly greater than or less than 0 are highlighted in red and blue, respectively, for the stimulation response. **C**, Comparison between data and simple model predictions. Top: all trials with sham stimulation and 100% noise. Bottom: varying image noise level or EVI independently. **D**, COD from models fit on broadband power at el-3 from main site stimulation. The image model yielded the highest COD, significantly greater than the simple model (*paired *t*-test, right-tailed P < 0.05). **E**, Bootstrapped means and 95% confidence intervals for image model responses (only means shown for the visual responses). **F**, Comparison between data and simple, image model predictions. Model outputs for electrodes 2 and 4 and control stimulation sites are presented in Figure S3.

Despite a prominent stimulation-induced broadband response, SPES in visual pathways did not modulate visual broadband responses as it did VEPs: the EVI model did not improve COD over the simple model (Figure 5A, D). Broadband responses were best described by the simple or image models. In subject 1, the simple model explained 48.5% of variance and was not significantly improved by any other model (Figure 5A). Thus, the single visual broadband response predicted by the simple model aligns well with the observed broadband responses, regardless of image noise level and EVI (Figure 5C). In subject 2, the image model explained significantly more variance (COD = 73.3%) than the simple model (Figure 5D), which means that predictions varying by image noise level better represented the observed data (Figure 5F). Increasing image noise was associated with longer sustained broadband elevation (Figure 5E). SPES at the control stimulation sites did not modulate visual broadband responses either, and it produced smaller stimulation-induced broadband responses than stimulating in visual pathways (Figure S3B).

### 3.4. Effects of SPES on visual processing outside early visual cortex

Here, we examine the effects of SPES on visual processing outside of the EVC to gauge the anatomical selectivity of stimulation effects. We analyzed data from measurement electrodes outside of the EVC that showed a significant broadband response to images in sham trials (subject 1: Figure S4, electrodes 5-10; subject 2: Figure S5, electrodes 11-13). These electrodes were located in LO1, the precentral sulcus and the inferior frontal gyrus.

SPES significantly modulated visual evoked potentials only at the two measured LO1 electrodes (Figure S4C, electrodes 5 and 6). Here, we observed a weaker but similar pattern of modulation as at EVC electrodes: stimulation with EVI = 0 or 100 ms yielded slightly larger VEPs than stimulation with EVI = 200 ms or sham stimulation. Notably, the modulation at electrode 6 was due to stimulation at the control site, rather than the main site. Overall, these weaker effects may reflect feedforward propagation from EVC. Stimulation did not affect the visual broadband responses at any electrode.

In general, for a given stimulation–measurement pair, SPES could have three alternative effects on either broadband or evoked potentials. First, it could produce no significant stimulation response (by bootstrapping, see supplemental methods). Second, it could produce a significant stimulation response but fail to modulate the visual response (i.e., simple or image model fit best). Third, it could produce a significant stimulation response *and* modulate the visual response (i.e., EVI or full model fit best). The two-way effects of SPES on evoked potentials vs. broadband, across all stimulation–measurement pairs, are presented in Table S1. This shows that SPES which modulated VEPs was also more likely to induce a stimulation broadband response (Table S1, third row), whereas SPES which failed to modulate VEPs tended to induce no broadband response (Table S1, second row). Thus, SPES modulation of VEPs appears to be correlated with its success in inducing a stimulation broadband response.

## 4. Discussion

In this study, we tested the modulatory potential of single pulse electrical stimulation (SPES) on visual evoked neuronal activity. Specifically, we quantified the effects of SPES in white matter tracts on evoked potentials and broadband changes in the visual system, both at rest and during a visual discrimination task. We found that visual white matter stimulation produced large amplitude evoked potentials in the early visual cortex (EVC), as well as transient broadband power increase followed by a longer decrease below baseline. During the visual discrimination task, SPES affected evoked potentials and broadband power in different ways: whereas it modulated the shape and amplitude of visual evoked potentials, it did not modulate visual broadband responses but produced stimulation broadband changes that directly added to them.

### 4.1. Single pulse electrical stimulation enhances visual evoked activity

Our results show that a single electrical pulse delivered in the visual pathways could enhance closely timed visual evoked potentials (VEPs). The timing of the single pulse with respect to the visual stimulus was important, as the VEP enhancement was only observed when stimulation was delivered briefly before or at visual onset (≤100ms). Interestingly, this time course is comparable to that of intracortical facilitation seen in paired pulse stimulation experiments with transcranial magnetic stimulation and intracranial stimulation^37^. Visual evoked potentials are thought to be driven by synchronous inputs from the thalamus and other cortical areas^24,25^. These results therefore suggest that visual white matter stimulation may induce a brief neuronal response in the visual cortex that facilitates those synchronous inputs.

### 4.2. Single pulse electrical stimulation affects total broadband activity

At early visual measurement electrodes, the stimulation-induced broadband response was characterized by a transient, several-fold increase in power followed by a sustained decrease. This pattern resembled our previous findings in the ventral temporal cortex when SPES was delivered to the hippocampus^18^, and is also consistent with previously reported effects of LGN stimulation on V1^17^.

The sustained broadband decrease from stimulation implies a relative suppression in total asynchronous neuronal firing in the visual cortex after the initial ∼100 ms of image onset. Note that unlike evoked potentials, broadband power has no meaningful polarity, and so its decrease from baseline does not mean the superposition of a distinct, *negative* signal. Rather, suppression of ongoing activity is a more likely explanation^17^. Mechanistically, the sustained stimulation-induced suppression could arise from refractory inhibition due to presynaptic neurotransmitter depletion, afterhyperpolarization, or postsynaptic receptor desensitization after the initial transient excitation^38–40^.

The stimulation-induced broadband response was simply additive to the visual broadband changes. A consequence of this addition is that differences in total broadband power are observed depending on the electrical-visual interval (EVI). When stimulation and visual onset are simultaneous, the two transient broadband peaks “stack” to yield greater total broadband power immediately after visual onset. But when stimulation and visual onset are offset by 200 ms, the visual broadband peak is counteracted by the suppressive phase of the stimulation broadband response, to yield lesser total broadband power immediately after visual onset. This difference might be meaningful if total population activity in the visual cortex influences downstream processing.

### 4.3. Visual modulation explained by added noise

While single pulse electrical stimulation had different effects on evoked potentials and broadband responses, our results showed that modulation of VEPs and a significant stimulation-induced broadband response often occurred together. This general pattern was seen across all visually responsive measurement electrodes, including those outside the EVC: SPES that modulated the VEP almost always produced a broadband response (Table S1, third row), while SPES that did not modulate the VEP mostly failed to produce a broadband response (Table S1, second row). This suggests that a shared underlying mechanism, like added noise, may explain both effects.

If a single electrical pulse adds noise to a downstream neuronal population, our data may be explained by the phenomenon of stochastic resonance^12,15^. In stochastic resonance, a small amount of random noise can boost periodic or event-related inputs above a neuronal firing threshold. The net effect is an increase in the synchrony and amplitude of signals at the neuronal population level. Too much noise, however, would drown the signal. Since broadband power is thought to reflect Poisson-distributed, noisy neuronal inputs^20^, the addition of more broadband power after visual onset by the stimulation-induced transient may be interpreted as the addition of random noise. And since evoked potentials are thought to reflect synchronous inputs, it follows that the VEPs after stimulation are amplified via stochastic resonance.

Added noise could be implemented at the pre- or post-synaptic level. At the presynaptic axon terminals of stimulated white matter tracts, SPES may cause a temporary buildup of Ca^2+^ that primes subsequent neurotransmitter release in response to visual stimuli^41–43^. Postsynaptically, additional excitatory NMDA receptors could be activated by glutamate spillover from the closely timed electrical and visual events^43,44^. Both potential mechanisms lead to larger excitatory postsynaptic potentials, which result in a larger VEP when summed across the neuronal population.

### 4.4. Alternative explanations for visual modulation

Single pulse transcranial magnetic stimulation has been shown to reset the phase of ongoing low frequency oscillations in scalp EEG^45^. If SPES can achieve a similar result, it could align specific phases of ongoing oscillations with the subsequent visual inputs, potentially modulating neuronal responses to those inputs^46^. Currently, this is not well explored in SPES, and testing this potential explanation may require careful analysis of phase relationships after SPES and immediately before VEPs.

The observed VEP facilitation might also be explained by rebound hyperactivity in visual cortex neurons subsequent to a more rapid, transient suppression; Kara et al.^47^ observed this phenomenon in cats when microstimulation was delivered to thalamic neurons with receptive fields close to measured V1 simple cells. However, an initial transient decrease in broadband power, not increase, would be expected immediately following stimulation. Thus, while rebound activity may be present in single cells, our population broadband measurements did not reveal such effects.

### 4.5. Single pulse electrical stimulation and behavioral threshold

Despite electrophysiologic evidence of visual facilitation, we observed inconsistent effects of stimulation on reaction time and no effect on response accuracy (see supplemental results, Figures S7-S8, and Tables S2-S4). This was expected and in line with other SPES experiments, since the charge delivered was titrated to be above the electrophysiologic response threshold^48^, but below the threshold of consistent behavioral detection. Stimulation thresholds for modulating evoked potentials, broadband activity, and behavior likely differ from one another and require careful titration. Higher amplitude stimulation may improve behavioral detection but also activate a larger volume of tissue, potentially obscuring the desired neurophysiology. For example, low current delivered to monkey MT has been shown to bias motion detection towards the direction encoded by stimulated neurons, but higher current simply impaired overall performance^49^.

Stimulating with pulse trains may also improve behavioral detection, like during clinical functional mapping of eloquent cortex^50^. However, different stimulation frequencies can have contrasting effects on neuronal firing rates, thus adding a parametric dimension to carefully consider^17,51^. Our goal is to first understand how the basic single pulse unit affects visual electrophysiology. Future research can then develop models linking these electrophysiological effects to behavioral outcomes from varying frequency and amplitude.

### 4.6. Limitations

The experiments in this study were conducted in two young male adult humans, which potentially limits the broader generalizability of our findings. iEEG electrodes were placed clinically and not optimized to be identical in both subjects, though diffusion MRI confirmed the anatomical similarity of the main stimulation and measurement electrodes with respect to the visual cortex and major white matter pathways (Figure 2). Across these subjects, stimulation likely simultaneously activated several major white matter tracts (e.g., ILF, IFOF, OR) and we cannot presently pinpoint which pathway(s) might be chiefly responsible for the effects on evoked potentials and broadband. Despite potential differences in electrode placement and white matter activated, we discovered similar evoked potential shapes, broadband responses, and time interval-dependent modulation in both subjects.

## 5. Conclusion

The experimental combination of SPES and visual inputs allowed us to uniquely observe how a single electrical pulse affects the neuronal processing of independent information. SPES in white matter tracts affected the visual cortex in a way consistent with other studies, showing a brief increase in population neuronal activity followed by longer lasting inhibition. These stimulation-induced broadband changes directly added to visual induced broadband changes. In contrast, visual evoked potentials, representative of synchronous visual inputs, were facilitated by SPES. These two complementary aspects of iEEG provide evidence for a shared underlying mechanism, e.g., that electrical stimulation adds neuronal noise. Combining electrical and visual inputs like this may serve well to elucidate the basis for even more complex neurostimulation paradigms on neuronal processing.

## Acknowledgements

We are grateful for the participation of the patients in this study, and for the assistance of Cindy Nelson, Karla Crockett, and other staff at Saint Mary Hospital, Mayo Clinic, Rochester, MN. Funding: Research reported in this publication was supported by the National Institutes of Health [R01 MH122258 to DH; R01 EY035533 to DH, KNK, and GAW; U01 NS128612 to KJM and GAW; T32 GM145408 to the Mayo Clinic MSTP] and by the American Epilepsy Society [937450 to HH].

## Data and code availability

The data that support the findings of this study will be available in Brain Imaging Data Structure (BIDS) format on OpenNeuro, upon manuscript publication. The code used to generate all results and figures will be available on GitHub.

## CRediT authorship contribution statement

**Harvey Huang:** Conceptualization, Methodology, Software, Validation, Formal Analysis, Investigation, Data Curation, Writing – Original Draft, Writing – Review & Editing, Visualization. **Kendrick N. Kay:** Conceptualization, Methodology, Validation, Formal Analysis, Writing – Review & Editing, Funding Acquisition. **Nicholas M. Gregg**: Investigation, Writing – Review & Editing, Supervision. **Gabriela Ojeda Valencia:** Data Curation, Writing – Review & Editing.

**Myung-Ho In**: Resources, Writing – Review & Editing. **Christoph Kapeller**: Methodology, Software, Writing – Review & Editing. **Yunhong Shu:** Resources, Writing – Review & Editing. **Gregory A. Worrell**: Resources, Supervision, Project Administration, Funding Acquisition. **Kai J. Miller:** Methodology, Resources, Writing – Review & Editing, Supervision, Funding Acquisition. **Dora Hermes:** Conceptualization, Methodology, Software, Validation, Formal Analysis, Investigation, Resources, Data Curation, Writing – Original Draft, Writing – Review & Editing, Visualization, Supervision, Project Administration, Funding Acquisition.

## Notes

### Competing Interest Statement

Christoph Kapeller reports a relationship with g.tec that includes: employment. Nicholas M. Gregg reports financial support was provided by Medtronic Inc. and a relationship with NeuroOne Inc. that includes: consulting or advisory. Gregory A. Worrell has patent licensed to Cadence Neuroscience Inc. and NeuroOne Inc. The other authors declare that they have no known competing financial interests or personal relationships that could have appeared to influence the work reported in this paper.

