## Supplementary materials for "Single pulse electrical stimulation in white matter modulates iEEG visual responses in human early visual cortex"

[1]Mayo Clinic Medical Scientist Training Program, Mayo Clinic, Rochester, MN; [2]Center for Magnetic Resonance Research, Department of Radiology, University of Minnesota, Minneapolis, MN; [3]Department of Neurology, Mayo Clinic, Rochester, MN; [4]Department of Physiology and Biomedical Engineering, Mayo Clinic, Rochester, MN; [5]Department of Radiology, Mayo Clinic, Rochester, MN; [6]g.tec medical engineering GmbH.; [7]Department of Neurologic Surgery, Mayo Clinic, Rochester, MN.

### Supplemental methods

#### Identification of visually responsive electrodes outside of early visual areas

Visually responsive measurement electrodes outside of early visual areas were functionally identified by their increased broadband power following visual onset during sham stimulation trials. Specifically, the power spectral density (PSD) was estimated using Welch's method for the 500 ms intervals before and after visual onset, for all sham trials at each electrode. A mean log broadband power was calculated for each interval by averaging the log power of 1-Hz frequency bins from 70-170 Hz, excluding the 116-124 Hz bins to avoid contamination from the 120 Hz line noise harmonic. For each electrode, an $R^2$ value was computed as the fraction of variance explained by the binary categorical variable, before vs. after visual onset, across all intervals. Electrodes were classified as visually responsive if they showed higher average log broadband power after visual onset and a statistically significant $R^2 > 0.1$ (FDR-corrected F-test, $P < 0.05$).

#### Diffusion MRI acquisition and preprocessing

Diffusion MRI (dMRI) scan was performed in both subjects on a 3T Prisma MRI scanner (Siemens Healthineers, Forchheim, Germany). A series with two volumes at b = 0-100 s/mm$^2$ and 60 directions at b = 1000 s/mm$^2$ were acquired with TR = 5100 ms; TE = 71 ms; 46 slices at 4 mm thickness (zero gap), field of view of 220 mm and acquisition matrix of 128 x 128.

The T1-weighted and dMRI images were preprocessed using QSIprep. The T1-weighted image was corrected for intensity non-uniformity[1] and then skull stripped using ANTS 2.3.1. Spatial normalization to the ICBM 152 Nonlinear Asymmetrical template version 2009c[2] was performed through nonlinear registration with antsRegistration[3], after brain-extraction of both the T1-weighted image volume and the template. Brain tissue segmentation was performed using FAST (FSL 6.0.3:b862cdd5[4]).

Any dMRI images with a b-value less than 100 s/mm$^2$ were treated as a b = 0 image. MP-PCA denoising was applied with a 5-voxel window[5], and B1 field inhomogeneity was corrected using dwibiascorrect (MRtrix3) with the N4 algorithm[1]. The mean intensity of the dMRI series was adjusted to match across b = 0 images from different dMRI scans. Head motion and eddy current were corrected using FSL's eddy (version 6.0.3:b862cdd5, q-space smoothing factor = 10, 5 iterations, 1000 voxels used to estimate hyperparameters[6]). Linear first and second level models were used to characterize eddy current-related spatial distortion and eddy's outlier replacement was run[7]. For outlier detection, data were grouped by slice, only including slices with at least 250 intracerebral voxels. Groups deviating by more than 4 standard deviations from the prediction value were replaced with imputed values. Final interpolation was performed using the jac method. The dMRI time-series were then resampled with 1.25 mm isotropic voxels in AC-PC space.

#### Confidence intervals for finite impulse response coefficients

Each FIR model was bootstrapped (1000 times for evoked potentials and 200 times for broadband responses) to calculate per-time point confidence intervals for stimulation and visual component coefficients. Resampling was done within each experimental condition to maintain balanced trial counts. In the EVI model, bootstrapped differences between pairs of visual responses were analyzed to locate time intervals showing significant EVI-dependent modulation: Consecutive time points spanning ≥20 ms were considered significantly different between pairs of EVI conditions (e.g., 0 ms vs. 100 ms) if the 99% confidence interval for the differences excluded 0 at all time points. For the analysis in section 3.4, a stimulation response was considered significant only if ≥20% of total time points within the first 500 ms post-stimulation had 95% confidence intervals that excluded 0.

#### Finite impulse response analysis of broadband responses

FIR analysis on broadband-transformed SPES+Visual data proceeded similarly as that on evoked potential data. Broadband power was downsampled to 200 Hz and temporally smoothed in log space with 50 ms time windows before analysis.

A key decision in broadband analysis is whether to fit models on power or log power, as addition in log power corresponds to multiplication in power space (Figure S6A), and both approaches have precedence in the literature[8,9]. Fitting on log power is statistically convenient because log power is approximately normally distributed whereas (raw) power is right-skewed; however, independent signals are theoretically additive in power when their phases are random[9]. To guide this decision, we compared the mean errors of image models fit on power vs. log power across all stimulation–measurement electrode pairs. Models estimated responses using least absolute deviation (LAD) regression, rather than least squares regression, because LAD regression targets the trial median – a quantity equivalent between power and log power up to the transformation itself. Trials were split into training (odd) and testing (even) for each experimental condition, and the training trials were used to fit image models on power and on log power (Figure S6B). We then calculated the mean absolute error (MAE) of power and log power model predictions, relative to the median observed testing trial for each experimental condition. We compared MAE between power and log power models across all 28 experimental conditions for all stimulation–measurement electrode pairs. On average, fitting on power yielded significantly lower MAE than fitting on log power (Figure S6C, paired *t*-test, $P < 0.05$). Thus, we moved forward using power for the broadband FIR analysis.

The same FIR models applied to evoked potentials were applied to the observed broadband power changes, with a few minor differences. Each model (simple, image, EVI, full) again followed the form $y = X\beta$, where y is a vector of observed broadband changes concatenated across all trials, X contains sequences of predictors corresponding to hypothesized broadband changes induced by stimulation and visual components, and β is the per-time point power of component responses to solve for. Instead of least squares regression used for evoked potentials, we continued to used LAD regression to solve for β, as it is more robust to the right-skewed distribution of broadband power. Each sequence of predictors in X began at 50 ms before stimulation or visual onset to account for temporal smoothing and smearing caused by forward-backward filtering in broadband preprocessing.

To maintain consistency with the evoked potential analysis, we again calculated COD on testing trials to compare the model performances. Before COD calculation, broadband predictions and observations were log-transformed to approximate normality and ensure the robustness of COD. As broadband changes appeared to persist longer than evoked potentials, we extended the time window of COD calculation to 1 s after visual onset. COD calculated up to 0.5 s, consistent with evoked potential results, are additionally presented in Figure S3C-D.

##### Evaluating effects of button press in finite impulse response models

To account for possible premotor or motor activity associated with button press, we added 2 additional sets of predictors to each best-fit FIR model, time-locked to button presses for each hand. These predictors were fit from -0.2 to 0.5 s around button press for evoked potentials and -0.25 to 0.5 s for broadband responses. In subject 1, these predictors also accounted for possible brain activity associated with the auditory feedback synchronized with button press. A significant increase in COD would indicate evoked potentials or broadband changes attributable to button press activity, but neither was observed.

### Supplemental results

#### SPES did not consistently impact reaction time or response accuracy

The main goal of the experiment was to test how a single electrical pulse modulates evoked and induced visual neuronal activity. However, it is valuable to assess whether a single electrical pulse influences perceptual measurements.

Subjects were asked to press a button to indicate whether noise masked images showed an *Elephant* or *Pizzas.* Mean reaction time differed significantly across the seven image conditions (subject 1: one-way ANOVA, $F(6, 641) = 7.71$, $P = 5.1*10^{-8}$; subject 2: one-way ANOVA, $F(6, 419) = 20.2$, $P = 8.4*10^{-21}$). As expected, the mean reaction time was significantly shorter for images with less noise (Figure S7A, D). To test whether reaction time was affected by stimulation, we fit a multivariate linear regression model, which adjusted for the run number, the trial order/timing, and the image category (Table S2, Figure S8). In subject 2, stimulation at the main site simultaneous with visual onset increased reaction time by 110 ms on average ($t(411) = 2.12$, $P = 0.035$), while stimulation at the control site 100 ms before visual onset decreased reaction time 88 ms on average ($t(411) = -2.19$, $P = 0.029$). However, these effects were not found in subject 1. Both subjects showed possible learning effects: subject 1 responded faster across trials within each run ($t(633) = -3.29$, $P = 0.0011$), and subject 2 responded faster with subsequent runs ($t(411) = -5.54$, $P = 5.3*10^{-8}$).

Response accuracy increased with less image noise, and it was significantly above chance level for 75%, 50%, and 0% noise images (Figure S7B, E). Whether images contained *Elephant* or *Pizzas* did not significantly affect accuracy for either subject (multivariate logistic regression, Table S3), so they were pooled to create psychometric (Weibull) functions of response accuracy for each stimulation site (Figure S7C, F). Stimulation at any site or with any EVI did not significantly influence response accuracy (Chi-Square tests of independence, Bonferroni-corrected $P > 0.05$ across 75%, 50%, and 0% noise levels), even after adjusting for run number, trial timing within runs, and image coherence (multivariate logistic regression, Table S4). The only robust predictor of response accuracy was image coherence (subject 1: $z = 9.20$, $P = 3.6*10^{-20}$; subject 2: $z = 7.56$, $P = 4.1*10^{-14}$).

#### Adjusted common average vs. bipolar re-referencing

In our evoked potential analysis, data were re-referenced by an adjusted common average that mitigated the inclusion of bias from other responsive channels[10]. Local re-referencing methods, such as bipolar or Laplacian, were avoided because they can strongly distort or attenuate meaningful evoked potentials between neighboring electrodes[10]. This can strongly confound interpretation if stimulation has no effect on the electrode of interest but evokes potentials at a neighboring electrode. However, highly focal signal features such as broadband changes might be better accentuated by local re-referencing[11]. Thus, we also tested broadband analysis after bipolar re-referencing (electrodes 1-2 and 3-4 in Figure S3).

Bipolar re-referencing reduced the peak amplitude of the stimulation broadband transients compared to adjusted common average re-referencing. This was likely attributable to better attenuation of sharp evoked potential peaks before filtering. This reduction was less pronounced in subject 1 than subject 2, and in neither case was the transient entirely eliminated. Therefore, the transients were unlikely to be filtering artifact alone.

Control stimulation in subject 1 modulated the visual induced broadband response when the data were bipolar re-referenced but not when adjusted common average re-referenced. (Figure S3B). This was the only case showing modulation of the visual broadband response. However, the increase in variance explained was marginal over the next best model, and the visual response time courses did not differ noticeably across stimulation conditions, so cautious interpretation is warranted. All other results were consistent regardless of reference choice.

### Supplemental figures

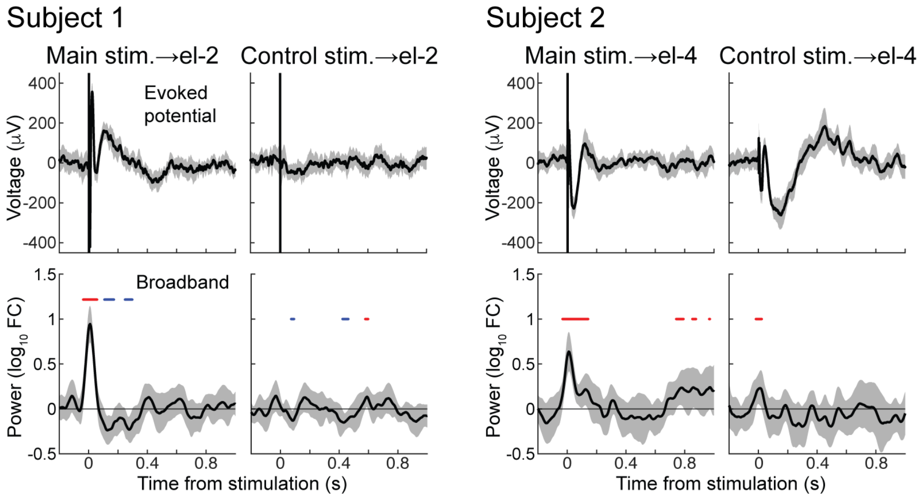


**Figure S1. BSEPs and induced broadband changes at measurement electrodes 2 and 4.** BSEPs (top) and induced broadband changes (bottom) recorded at el-2 in subject 1 and el-4 in subject 2 from main and control stimulation sites in each subject. Shaded intervals depict 95% confidence interval of the mean. Time points with mean broadband significantly greater than or less than 0 are highlighted in red and blue, respectively (one-sample *t*-test, $P < 0.05$).

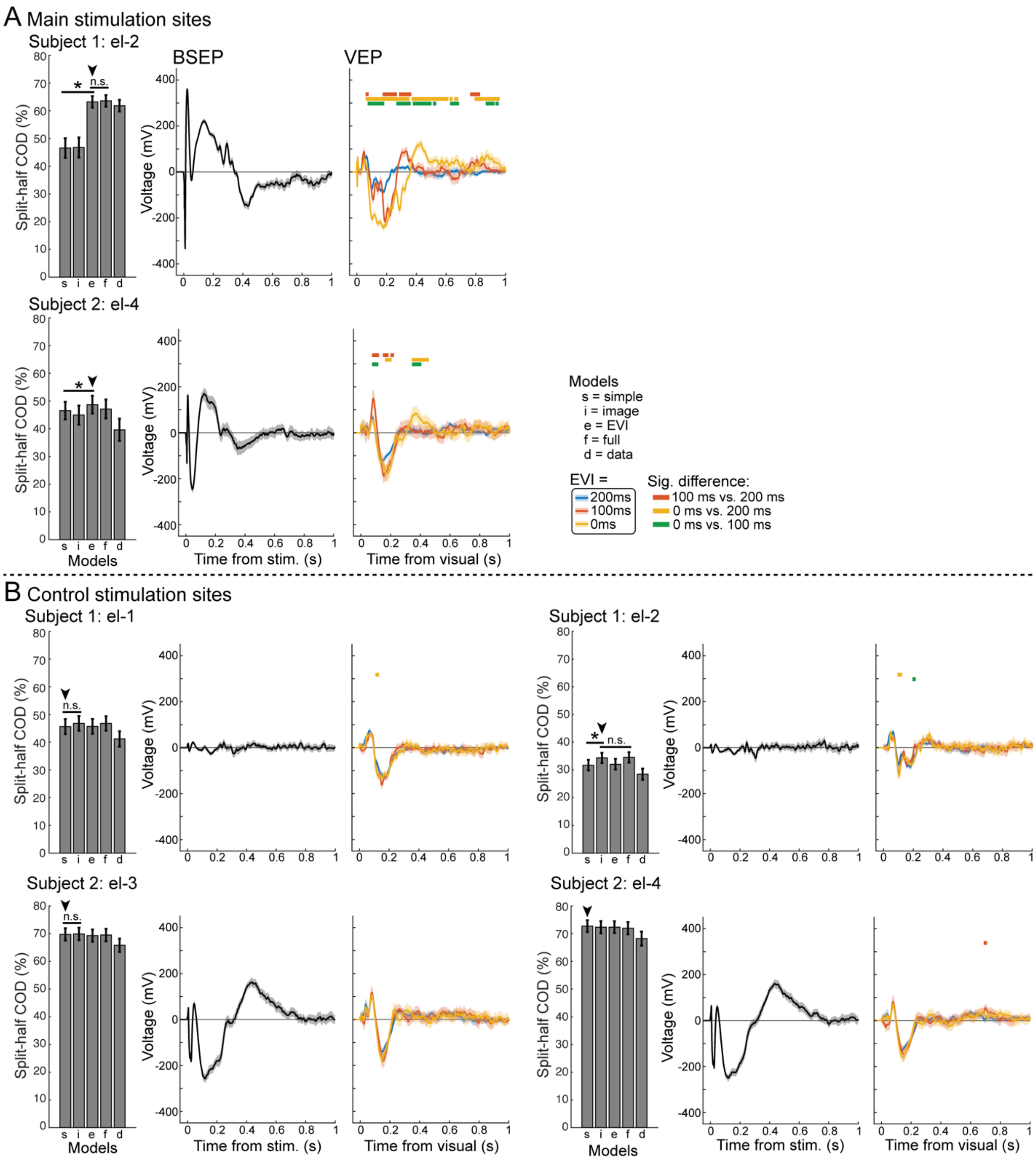


**Figure S2. Split-half COD and EVI model responses on evoked potential data for all other early visual measurement electrodes and control stimulation sites. A,** Results for main stimulation sites to el-2 (subject 1) and el-4 (subject 2). **B,** Results for control stimulation sites to all early visual measurement electrodes. Statistical comparisons for COD indicate paired *t*-test at right-tailed $\alpha = 0.05$ for the full model > EVI/image models, or for EVI/image models > the simple model. Arrowheads indicate the best model. Shaded intervals show 95% confidence intervals of the mean, and significantly different time intervals between pairs of EVI conditions are labeled with colored bars above (bootstrapped differences, $P < 0.01$).

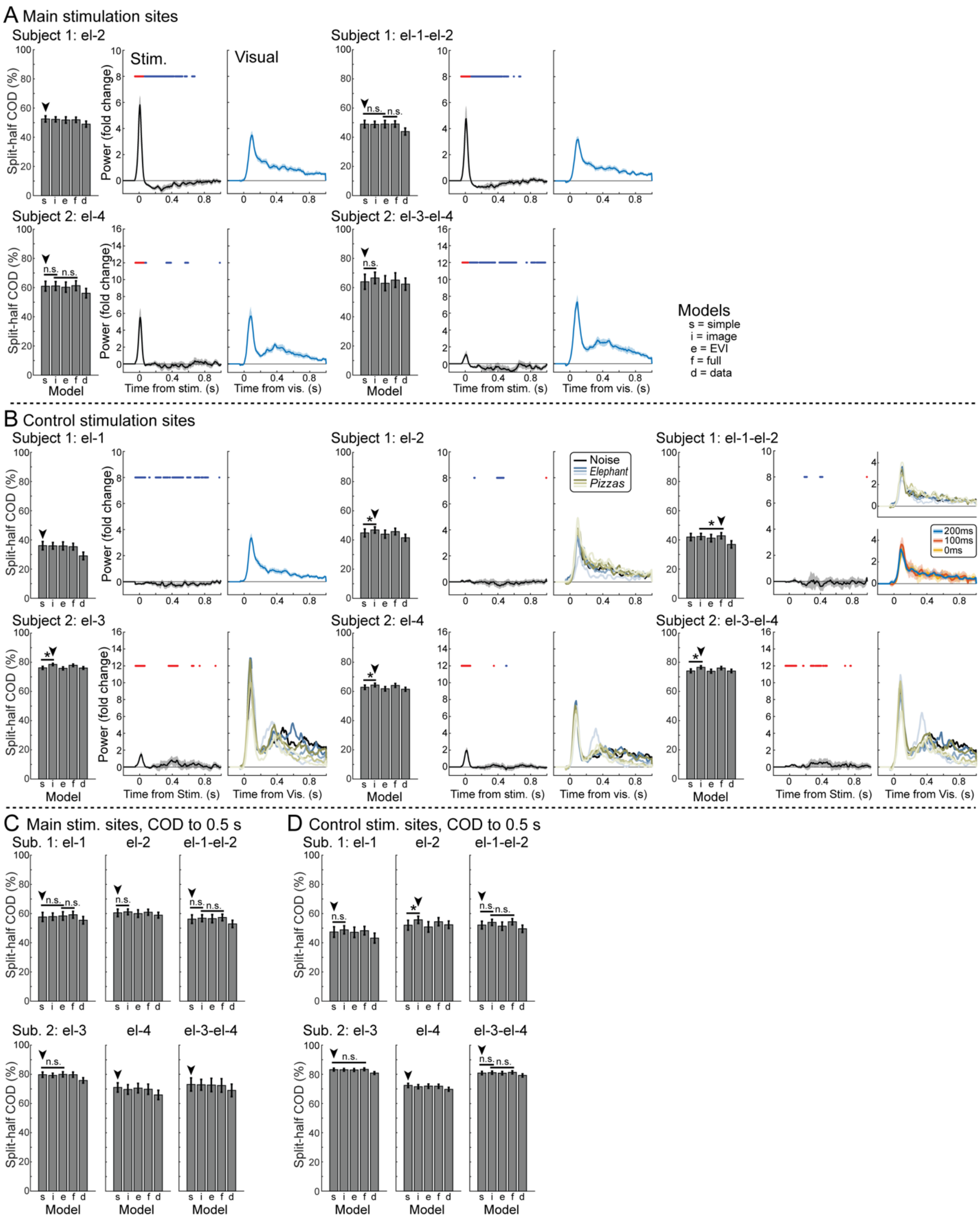


**Figure S3. Split-half COD and best-fit FIR model on broadband power for all other early visual measurement electrodes and control stimulation sites. A,** Results for main stimulation sites to el-2, el-4, and bipolar re-referenced electrode pairs, COD calculated up to 1 s after visual onset. **B,** Results for control stimulation sites to early visual measurement electrodes and their bipolar re-referenced pairs, COD calculated up to 1 s after visual onset. Statistical comparisons for COD indicate paired *t*-test at right-tailed $\alpha = 0.05$ for the full model > EVI/image models, or for EVI/image models > the simple model. Arrowheads indicate the best model. Shaded intervals show 95% confidence intervals of the mean for the best model (omitted for image models). Time points significantly greater than or less than 0 are highlighted in red and blue, respectively, for stimulation responses. Lighter colors in image model responses denote lower noise (75%, 50%, 0%). **C, D,** COD calculated up to 0.5 s after visual onset.

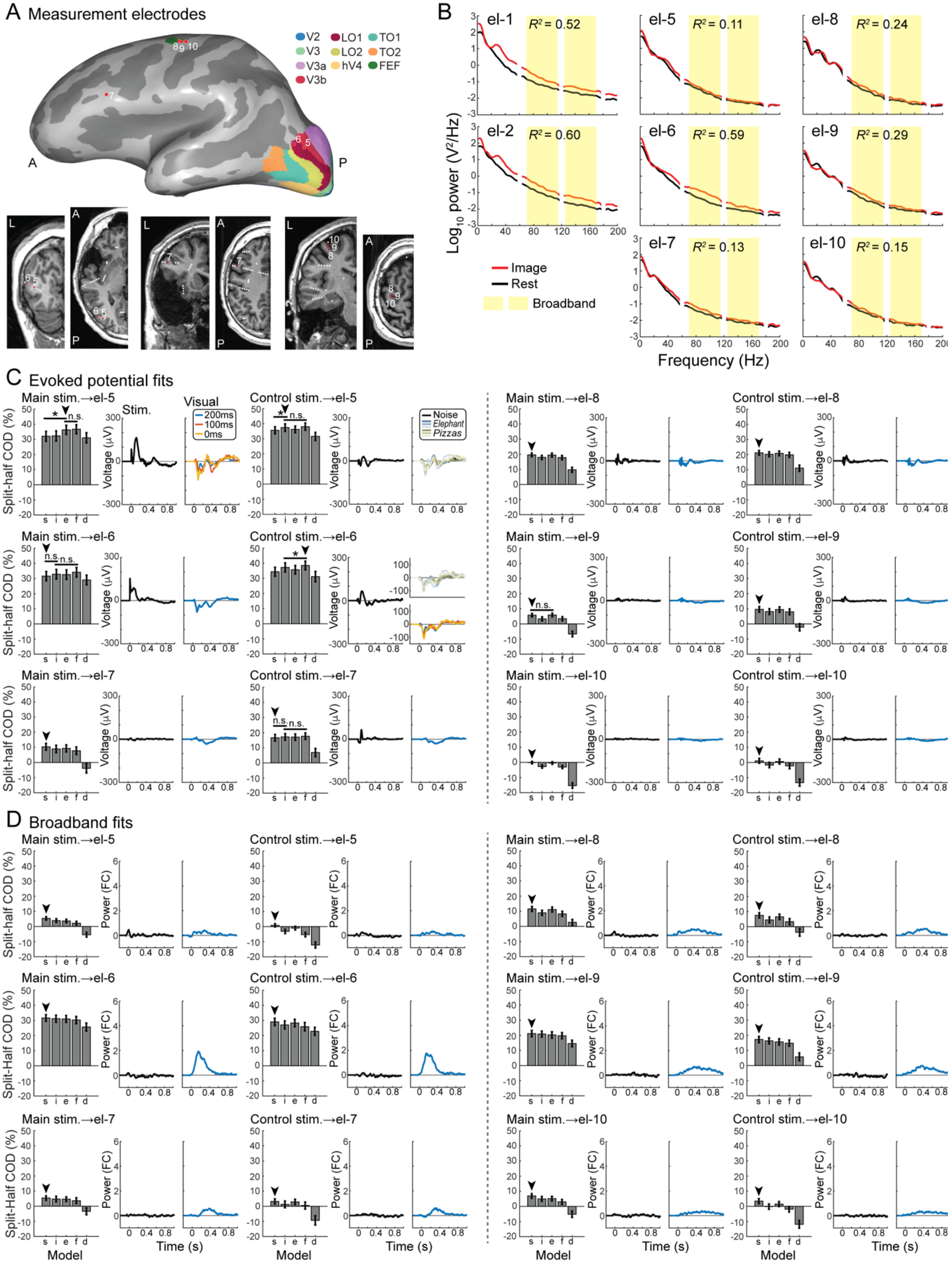

A Measurement electrodes
V2 V3 V3a V3b LO1 LO2 hV4 TO1 TO2 FEF
B
el-1 $R^2$ = 0.52
el-2 $R^2$ = 0.60
el-5 $R^2$ = 0.11
el-6 $R^2$ = 0.59
el-7 $R^2$ = 0.13
el-8 $R^2$ = 0.24
el-9 $R^2$ = 0.29
el-10 $R^2$ = 0.15
Log10 power (V²/Hz)
Frequency (Hz)
Image
Rest
Broadband
C Evoked potential fits
Main stim.→el-5
Control stim.→el-5
Main stim.→el-8
Control stim.→el-8
Main stim.→el-6
Control stim.→el-6
Main stim.→el-9
Control stim.→el-9
Main stim.→el-7
Control stim.→el-7
Main stim.→el-10
Control stim.→el-10
Stim.
Visual
200ms
100ms
0ms
Noise
Elephant
Pizzas
Split-half COD (%)
Voltage (μV)
D Broadband fits
Power (FC)
Model
Time (s)

**Figure S4. FIR model fits on evoked potential and broadband data at non-EVC visually responsive electrodes in subject 1. A,** Visually responsive measurement electrodes outside the EVC in subject 1 (5-10) visualized on the subject's inflated pial surface (top) and on subject coronal and axial T1-weighted MRI slices in red (bottom). Other electrodes within 4 mm of each slice are also plotted in white. **B,** Mean power spectral density plots for each visually responsive electrode across sham trials, before (rest) and after (image) visual onset. $R^2$ is the variance of broadband log power explained by condition, image or rest. **C,** COD and best model responses for evoked potentials at non-EVC recording electrodes. s = simple, i = image, e = EVI, f = full, d = data. **D,** COD (calculated up to 1 s after visual onset) and best model responses for broadband power at non-EVC electrodes. Statistical comparisons indicate paired *t*-test at right-tailed $\alpha$ = 0.05 for the full model > EVI/image models, or for EVI/image models > the simple model. Arrowheads indicate the best model.

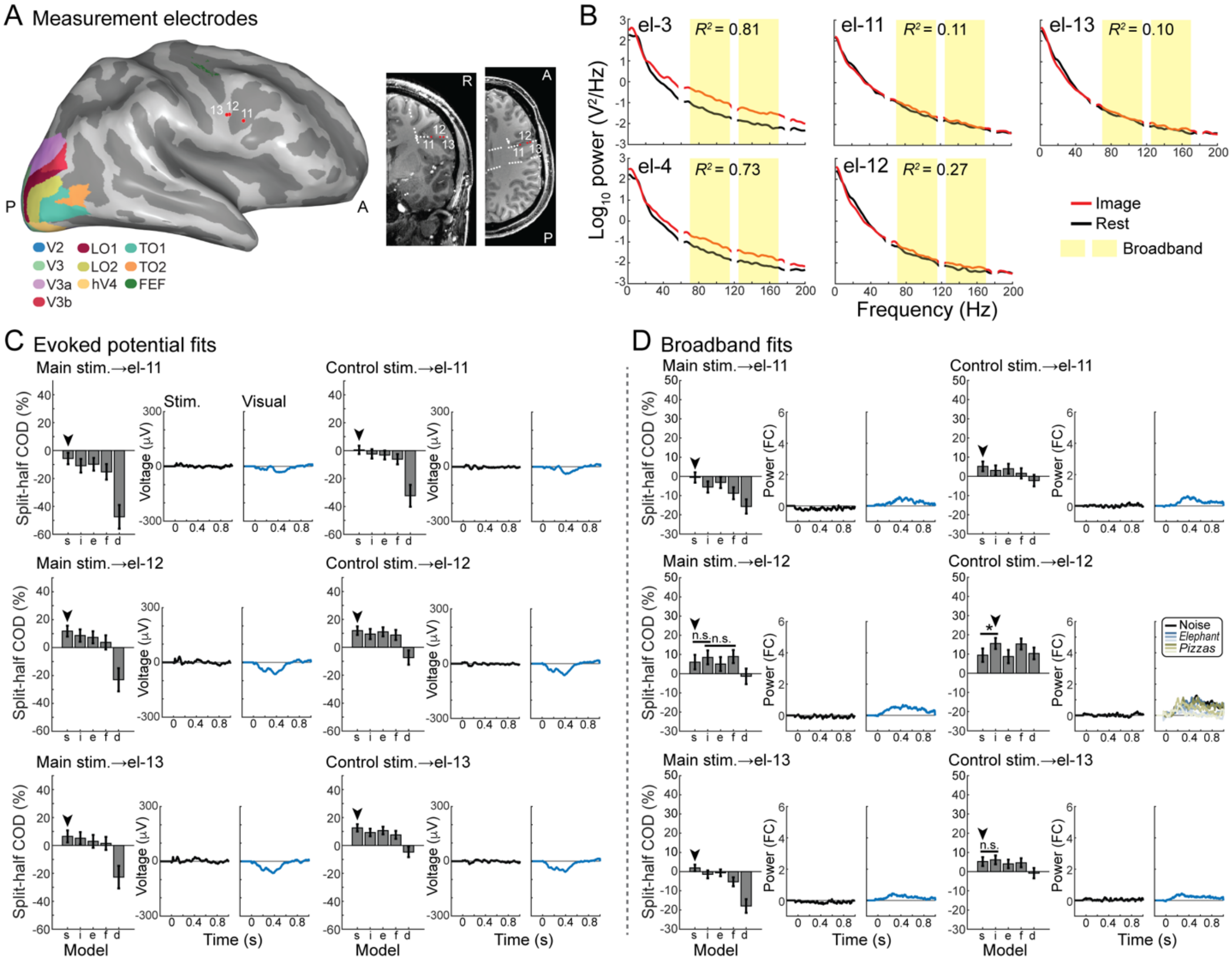


**Figure S5. FIR model fits on evoked potential and broadband data at non-EVC visually responsive electrodes in subject 2. A,** Visually responsive measurement electrodes outside the EVC in subject 2 (11-13) visualized on the subject's inflated pial surface (left) and on subject coronal and axial T1-weighted MRI slices in red (right). Other electrodes within 4 mm of each slice are also plotted in white. **B,** Mean power spectral density plots for each visually responsive electrode across sham trials, before (rest) and after (image) visual onset. $R^2$ is the variance of broadband log power explained by condition, image or rest. **C,** COD and best model responses for evoked potentials at non-EVC recording electrodes. s = simple, i = image, e = EVI, f = full, d = data. **D,** COD (calculated up to 1 s after visual onset) and best model responses for broadband power at non-EVC electrodes. Statistical comparisons indicate paired *t*-test at right-tailed $\alpha$ = 0.05 for the full model > EVI/image models, or for EVI/image models > the simple model. Arrowheads indicate the best model.

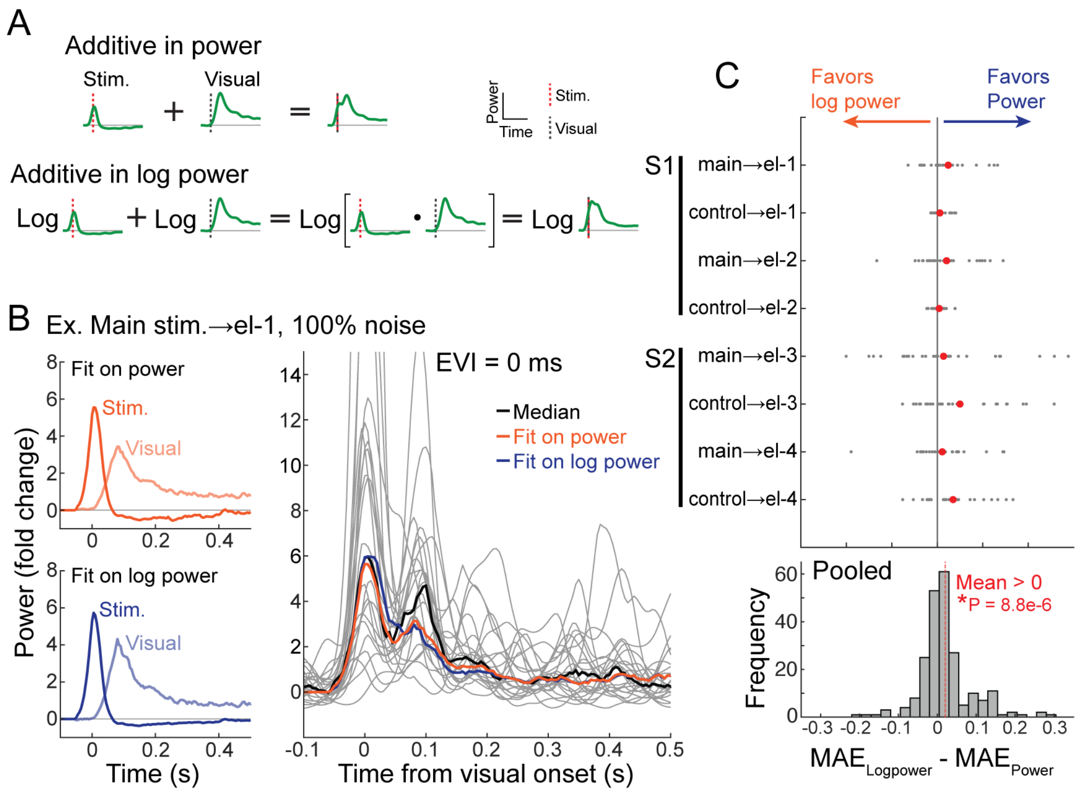


**Figure S6. FIR models fit better on broadband power than on broadband log power. A,** An FIR model fit on log power is mathematically equivalent to a multiplicative model fit on power. **B,** (Left) image model responses fit on broadband power vs. broadband log power, for an example stimulation-measurement electrode pair and one image condition. (Right) Comparison between data and the predictions from power and log power models. **C,** (Top) Difference in split-half mean absolute error (MAE) between log power and power image models for each stimulation-measurement electrode pair. Gray dots are single experimental conditions, and red circle is mean across experimental conditions. (Bottom) Distribution of differences in split-half MAE, pooled across all stimulation-measurement electrode pairs. Mean MAE is greater for the log power model fits compared to the power model fits.

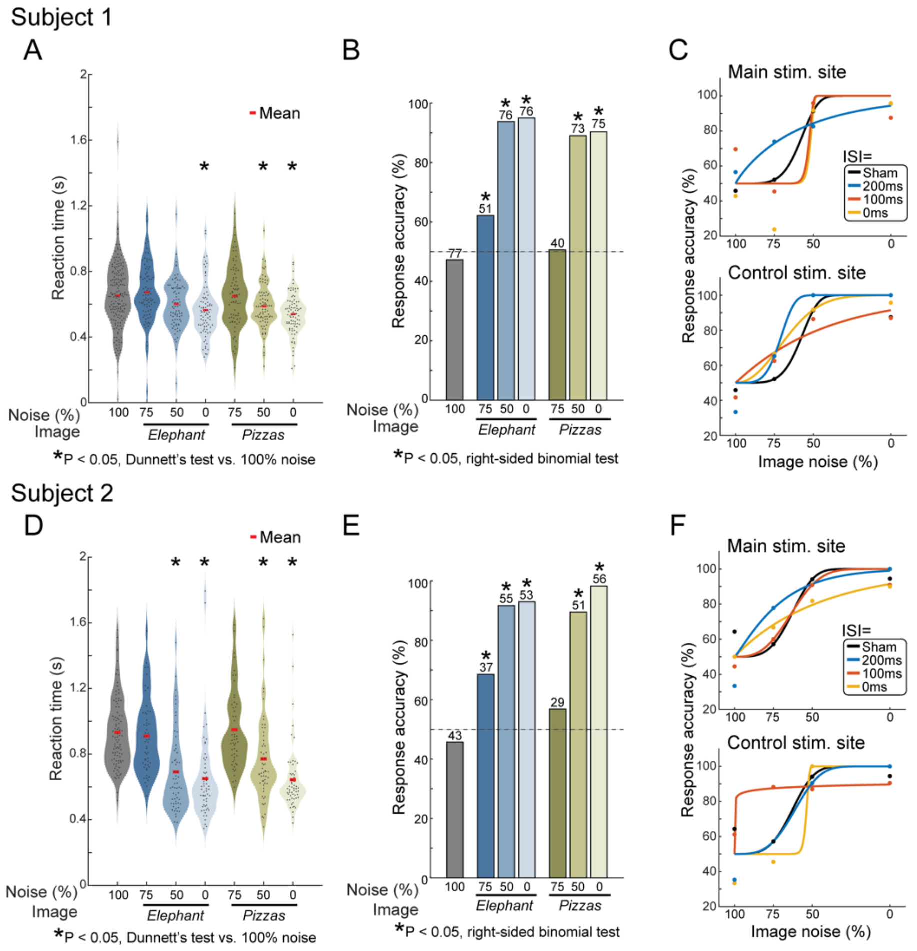


**Figure S7. Reaction time and response accuracy improve with less image noise. A,** Subject 1 response time for all trials, by image condition. **B,** Subject 1 response accuracy for each image condition. The number above each bar indicates the number of hits for that image condition. Chance level (50%) is shown with the gray dashed line. **C,** Response accuracy as a Weibull function of image noise level for each stimulation site. **D-F** show analogous results in subject 2 as in A-C.

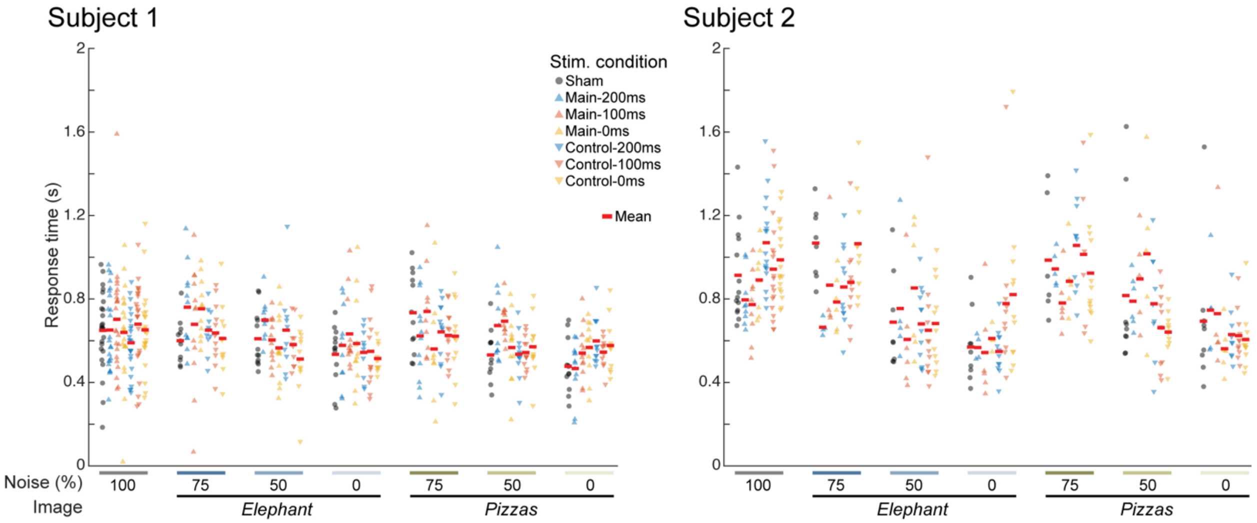


**Figure S8. Reaction times subdivided by image and stimulation condition.** Expansion of distributions in Figure S7A, D, by stimulation condition. Each point is a single trial.

#### Supplemental tables

**Table S1. Modulatory relationship between stimulation and visual responses for all stimulation→measurement electrode pairs[a]**

| EP \ Broadband | -SPES, -mod. | +SPES, -mod. | +SPES, +mod. |
|---|---|---|---|
| -SPES, -mod. | Control→el-11<br>Main→el-12<br>Control→el-12 | Main→el-11<br>Control→el-13 | |
| +SPES, -mod. | **Control→el-2**<br>**Control→el-4**<br>Main→el-7<br>Control→el-7<br>Main→el-8<br>Control→el-8<br>Main→el-9<br>Control→el-9<br>Main→el-10<br>Control→el-10 | **Control→el-1**<br>**Control→el-3**<br>Control→el-5<br>Main→el-6<br>Main→el-13 | |
| +SPES, +mod. | Control→el-6 | **Main→el-1**<br>**Main→el-3**<br>**Main→el-2**<br>**Main→el-4**<br>Main→el-5 | |

[a]Rows indicate the modulatory relationship between BSEPs and VEPs, and columns indicate the relationship between stimulation and visual induced broadband changes. "-SPES, -mod." = Stimulation did not produce a significant response and did not modulate visual response; "+SPES, -mod." = stimulation produced a response but did not modulate the visual response; "+SPES, +mod." = stimulation modulated the visual response (best-fit model was EVI or full). EVC electrodes are bolded, and main stimulation-measurement electrode pairs shown in Figures 4 and 5 are additionally in red.

#### Table S2. Multivariate linear regression of reaction time[b]

| | Coefficient | 95% CI | Test Statistic | P-value |
|---|---|---|---|---|
| **Subject 1** | | | | |
| Intercept | 0.71 | [0.65, 0.76] | $t(633) = 24.3$ | 7.2e-93* |
| Run number | -0.018 | [-0.039, 0.0027] | $t(633) = -1.71$ | 0.088 |
| Trial onset time (s) | -2.1e-4 | [-3.3e-4, -8.3e-5] | $t(633) = -3.29$ | 0.0011* |
| Main-200ms | 0.018 | [-0.035, 0.070] | $t(633) = 0.669$ | 0.50 |
| Main-100ms | 0.045 | [-0.0072, 0.098] | $t(633) = 1.69$ | 0.091 |
| Main-0ms | -0.0064 | [-0.059, 0.046] | $t(633) = -0.238$ | 0.81 |
| Control-200ms | 0.013 | [-0.040, 0.065] | $t(633) = 0.476$ | 0.64 |
| Control-100ms | 0.018 | [-0.034, 0.070] | $t(633) = 0.669$ | 0.50 |
| Control-0ms | 0.0089 | [-0.043, 0.061] | $t(633) = 0.337$ | 0.74 |
| *Elephant*-75% noise | 0.015 | [-0.030, 0.060] | $t(633) = 0.655$ | 0.51 |
| *Elephant*-50% noise | -0.053 | [-0.097, -0.0077] | $t(633) = -2.30$ | 0.022* |
| *Elephant*-0% noise | -0.090 | [-0.13, -0.045] | $t(633) = -3.92$ | 9.7e-5* |
| *Pizzas*-75% noise | -0.0039 | [-0.049, 0.041] | $t(633) = -0.172$ | 0.86 |
| *Pizzas*-50% noise | -0.063 | [-0.11, -0.018] | $t(633) = -2.77$ | 0.0057* |
| *Pizzas*-0% noise | -0.11 | [-0.16, -0.069] | $t(633) = -5.02$ | 6.6e-7* |
| **Subject 2** | | | | |
| Intercept | 1.0 | [0.95, 1.1] | $t(411) = 22.7$ | 1.1e-74* |
| Run number | -0.12 | [-0.17, -0.080] | $t(411) = -5.54$ | 5.3e-8* |
| Trial onset time (s) | 1.9e-4 | [-2.0e-5, 4.0e-4] | $t(411) = 1.78$ | 0.076 |
| Main-200ms | 0.065 | [-0.038, 0.17] | $t(411) = 1.24$ | 0.22 |
| Main-100ms | 0.044 | [-0.057, 0.15] | $t(411) = 0.861$ | 0.39 |
| Main-0ms | 0.11 | [0.0079, 0.21] | $t(411) = 2.12$ | 0.035* |
| Control-200ms | -0.065 | [-0.14, 0.014] | $t(411) = -1.62$ | 0.11 |
| Control-100ms | -0.088 | [-0.17, -0.0090] | $t(411) = -2.19$ | 0.029* |
| Control-0ms | -0.048 | [-0.13, 0.030] | $t(411) = -1.21$ | 0.23 |
| *Elephant*-75% noise | -0.026 | [-0.10, 0.051] | $t(411) = -0.671$ | 0.50 |
| *Elephant*-50% noise | -0.25 | [-0.32, -0.17] | $t(411) = -6.58$ | 1.4e-10* |
| *Elephant*-0% noise | -0.29 | [-0.36, -0.21] | $t(411) = -7.39$ | 8.1e-13* |
| *Pizzas*-75% noise | 0.016 | [-0.062, 0.095] | $t(411) = 0.407$ | 0.68 |
| *Pizzas*-50% noise | -0.17 | [-0.25, -0.094] | $t(411) = -4.41$ | 1.3e-5* |
| *Pizzas*-0% noise | -0.30 | [-0.37, -0.22] | $t(411) = -7.64$ | 1.6e-13* |

[b]Multivariate linear regression of response time for each subject. The intercept term represents the expected response time in sham stimulation trials when the run number is 1, the trial onset is at 0 s, and the image is 100% noise.

**Table S3. Multivariate logistic regression of response accuracy with image coherence, natural scene contained, and their interaction[c]**

| | Ln OR | 95% CI | Test Statistic | P-value |
|---|---|---|---|---|
| **Subject 1** | | | | |
| Image coherence (%) | 0.027 | [0.0181, 0.0354] | *z* = 6.06 | 1.3e-9* |
| Scene: *Elephant* vs. *Pizzas* | -0.18 | [-0.71, 0.35] | *z* = -0.667 | 0.51 |
| Coherence:*Elephant* | 0.014 | [-6.3e-4, 0.028] | *z* = 1.88 | 0.061 |
| **Subject 2** | | | | |
| Image coherence (%) | 0.046 | [0.030, 0.061] | *z* = 5.82 | 6.1e-9* |
| Scene: *Elephant* vs. *Pizzas* | 0.53 | [-0.17, 1.2] | *z* = 1.49 | 0.14 |
| Coherence:*Elephant* | -0.014 | [-0.034, 0.0062] | *z* = -1.35 | 0.18 |

[c]Multivariate logistic regression to determine whether image coherence (100% – image noise), original natural scene contained, or both are predictive of response accuracy.

**Table S4. Multivariate logistic regression of response accuracy with image coherence, run number, onset, and stimulation condition[d]**

| | Ln OR | 95% CI | Test Statistic | P-value |
|---|---|---|---|---|
| **Subject 1** | | | | |
| Intercept | -0.63 | [-1.4, 0.15] | *z* = -1.58 | 0.11 |
| Run number | 0.19 | [-0.12, 0.49] | *z* = 1.21 | 0.23 |
| Trial onset time (s) | 2.4e-6 | [-0.0018, 0.0018] | *z* = 0.0026 | 1.00 |
| Img. coherence (%) | 0.033 | [0.026, 0.040] | *z* = 9.20 | 3.6e-20* |
| Main-200ms | 0.68 | [-0.10, 1.5] | *z* = 1.71 | 0.088 |
| Main-100ms | 0.53 | [-0.24, 1.3] | *z* = 1.35 | 0.18 |
| Main-0ms | -0.094 | [-0.84, 0.65] | *z* = -0.246 | 0.81 |
| Control-200ms | 0.12 | [-0.64, 0.89] | *z* = 0.314 | 0.75 |
| Control-100ms | -0.18 | [-0.93, 0.58] | *z* = -0.456 | 0.65 |
| Control-0ms | 0.10 | [-0.66, 0.86] | *z* = 0.261 | 0.79 |
| **Subject 2** | | | | |
| Intercept | -0.046 | [-1.0, 0.92] | *z* = -0.0936 | 0.93 |
| Run number | -0.072 | [-0.59, 0.44] | *z* = -0.273 | 0.78 |
| Trial onset time (s) | 8.5e-4 | [-0.0016, 0.0033] | *z* = 0.686 | 0.49 |
| Img. coherence (%) | 0.039 | [0.029, 0.049] | *z* = 7.56 | 4.1e-14* |
| Main-200ms | -0.049 | [-1.2, 1.1] | *z* = -0.0801 | 0.94 |
| Main-100ms | -0.30 | [-1.5, 0.86] | *z* = -0.506 | 0.61 |
| Main-0ms | -0.31 | [-1.5, 0.84] | *z* = -0.527 | 0.60 |
| Control-200ms | -0.46 | [-1.4, 0.46] | *z* = -0.983 | 0.33 |
| Control-100ms | 0.21 | [-0.75, 1.2] | *z* = 0.427 | 0.67 |
| Control-0ms | -0.51 | [-1.4, 0.38] | *z* = -1.12 | 0.26 |

[d]Multivariate logistic regression of response accuracy for each subject. The intercept term represents the expected response accuracy in sham stimulation trials when the run number is 1, the trial onset is at 0 s, and the image coherence is 0% (100% noise).
